# A coarse-grained model for disordered and multi-domain proteins

**DOI:** 10.1101/2024.02.03.578735

**Authors:** Fan Cao, Sören von Bülow, Giulio Tesei, Kresten Lindorff-Larsen

## Abstract

Many proteins contain more than one folded domain, and such modular multi-domain proteins help expand the functional repertoire of proteins. Because of their larger size and often substantial dynamics, it may be difficult to characterize the conformational ensembles of multi-domain proteins by simulations. Here, we present a coarse-grained model for multi-domain proteins that is both fast and provides an accurate description of the global conformational properties in solution. We show that the accuracy of a one-bead-per-residue coarse-grained model depends on how the interaction sites in the folded domains are represented. Specifically, we find excessive domain-domain interactions if the interaction sites are located at the position of the C_*α*_ atoms. We also show that if the interaction sites are located at the centre of mass of the residue, we obtain good agreement between simulations and experiments across a wide range of proteins. We then optimize our previously described CALVADOS model using this centre-of-mass representation, and validate the resulting model using independent data. Finally, we use our revised model to simulate phase separation of both disordered and multi-domain proteins, and to examine how the stability of folded domains may differ between the dilute and dense phases. Our results provide a starting point for understanding interactions between folded and disordered regions in proteins, and how these regions affect the propensity of proteins to self-associate and undergo phase separation.

## Introduction

Multi-domain proteins (MDPs) consist of more than one folded domain that are often connected by linkers or longer intrinsically disordered regions (IDRs), and make up a large fraction (around 50%) of the proteomes in eukaryotic and prokaryotic organisms (***Han et al., 2007; Van Der Lee et al., 2014***). Like intrinsically disordered proteins (IDPs), MDPs can display large-amplitude motions that may play prominent roles in biomolecular functions like signalling, catalysis and regulation (***Mackereth and Sattler, 2012; Van Der Lee et al., 2014; Delaforge et al., 2016; Bondos et al., 2021***).

The biological functions of MDPs depend both on the properties of the folded domains and the disordered regions, and so characterizing the conformational ensembles can be key to understanding how these proteins function. In many cases, the folded and disordered regions are studied separately, but the folded domains might affect the conformational properties of the disordered regions (***Mittal et al., 2018; Taneja and Holehouse, 2021***) and the disordered regions may also affect the properties of the folded domains (***Yu and Sukenik, 2023***). For example, there is a complex interplay between the folded and disordered regions in the RNA-binding protein hnRNPA1, that affects its conformational ensemble in solution and its propensity to undergo phase separation (***Martin et al., 2021b***). However, describing the conformational ensembles of MDPs in solution generally requires a combination of biophysical experiments and molecular dynamics (MD) simulations (***Thomasen and Lindorff-Larsen, 2022***).

All-atom MD simulations have been used to generate conformational ensembles of IDPs and MDPs and to study intra- and inter-domain interactions (***Zheng et al., 2020; Sekiyama et al., 2022***). Such simulations, however, are often limited by the large system sizes and long time scales which limit efficient sampling of these dynamic proteins. Coarse-grained (CG) models may increase the sampling efficiency by reducing the number of particles in the simulation systems (***Neri et al., 2005; Monticelli et al., 2008; Bereau and Deserno, 2009; Gopal et al., 2010***). The accuracy, transferability, and efficiency of such models, however, depend on the degree of coarse-graining and the parameterization strategy (***Heo and Feig, 2024***). One commonly used model is the Martini force field, which uses a four-to-one mapping scheme with explicit solvent (***Souza et al., 2021***). Different versions of Martini have been modified to produce improved ensembles of IDPs and MDPs (***Benayad et al., 2020; Thomasen et al., 2022, 2023***). For IDPs, there has in the last years been extensive work using even coarser models where each amino acid residue is represented by a single bead. The interaction sites are generally located at the C_*α*_ position and separated by bonds that are 0.38 nm long, and we therefore here term these C_*α*_ models. Several related models rely on a similar functional form to the HPS model introduced by ***Dignon et al***. (***2018***) and may include bonded terms, an Ashbaugh-Hatch potential (***Ashbaugh and Hatch, 2008***) for shorter-range interactions and a Debye-Hückel electrostatic screening potential. Such models have for example been used to study the conformational ensembles and interactions within and between IDPs (***Dignon et al., 2018; Joseph et al., 2021; Regy et al., 2021; Dannenhoffer-Lafage and Best, 2021; Wessén et al., 2022; Tesei and Lindorff-Larsen, 2023; Valdes-Garcia et al., 2023***).

Coarse-grained models developed for IDPs do not represent the stability of folded proteins well, because the finely balanced energy contributions from individual backbone and side-chain interactions are not captured by the reduced representation. As a consequence, additional (often harmonic) restraints are applied to maintain the folded configurations in folded proteins and MDPs (***Souza et al., 2021; Borges-Araújo et al., 2023***). Even when applying such restraints to models developed for IDPs, extra attention needs to be paid to interactions related to folded domains since it is still unclear whether the models are fully transferable to MDPs. In particular, C_*α*_-based one-bead-per-residue mappings do not account for the specific orientations of side chains in folded proteins (***Kolinski and Skolnick, 1998***). For example, hydrophobic residues, whose side chains are ‘tucked away’ in the hydrophobic core of the protein, may be exposed at the surface of the protein in a C_*α*_ based representation. One approach to help overcome this problem is to use a different or scaled set of force field parameters for interactions that involve folded regions (***Kim and Hummer, 2008; Dignon et al., 2018; Krainer et al., 2021***). Another possible solution is the introduction of more terms in the energy function to better describe long-range interactions (***Li et al., 2012; Tan et al., 2023***) or to introduce anisotropic interactions (***Sieradzan et al., 2022***).

As an alternative, other coarse-grained models represent a residue by more than one bead to represent backbone side chain orientations and interactions (***Pappu et al., 1996; Hyeon et al., 2006; Maity et al., 2022; Zhang et al., 2022; Sieradzan et al., 2022; Mugnai et al., 2023; Zhang et al., 2023; Yamada et al., 2023***). In some of these models, one bead is placed at C_*α*_ and the other one is at the centre of mass (COM) of side chain atoms. In this way, side chain interactions can be explicitly taken into account, improving the simulated dynamical behaviour of folded protein simulations and model transferability. In previous studies, this strategy has been used to study conformational ensembles of IDPs or unfolding pathways of proteins (***Hyeon et al., 2006; Mugnai et al., 2023***). While effective, using multi-bead-per-residue models increases the time to sample configurations in simulations, and requires the determination of a larger number of force field parameters.

We have previously developed and applied an automated procedure to optimize the ‘stickiness’ parameters (*λ*) in a one-bead-per-residue model by improving the agreement with experimental small-angle X-ray scattering (SAXS) and paramagnetic relaxation enhancement (PRE) nuclear magnetic resonance (NMR) data for a large set of IDPs (***Norgaard et al., 2008; Tesei et al., 2021b; Tesei and Lindorff-Larsen, 2023***). The most recent CALVA-DOS (Coarse-graining Approach to Liquid-liquid phase separation Via an Automated Data-driven Optimisation Scheme) model (CALVADOS 2) was further tuned to describe phase behaviour of multi-chain conformational ensembles of IDPs from simulations by reducing the range of non-ionic interactions (***Tesei and Lindorff-Larsen, 2023***).

Here, we explore the use of the CALVADOS model for simulations of MDPs. We find that when the CAL-VADOS 2 parameters are used in simulations of MDPs with interaction sites at the C_*α*_ positions, the resulting structures in some cases show excessive interactions between the folded domains, leading to compact ensembles that do not agree with SAXS data. To remedy this problem, we describe a strategy where interaction sites in folded regions are located at the COM of the residue, and show that simulations with this model result in substantially improved agreement with experiments. We optimize the parameters in CALVADOS using the COM representation to derive a refined set of CALVADOS parameters (CALVADOS 3). When we combine the COM representation of folded domains with harmonic restraints between residues in the folded domains and the CALVADOS 3 parameters we obtain good agreement with experimental data on single-chain properties of MDPs and IDPs. Finally, we show how this model may be used to study the interactions between folded and disordered regions in proteins that undergo phase separation, and how the stability of folded domains might change during phase separation.

## Results

### A modified representation improves accuracy for multi-domain proteins

We first evaluated the accuracy of the original CALVADOS 2 model for simulations of MDPs. We therefore used the CALVADOS 2 parameters (***Tesei and Lindorff-Larsen, 2023***) and a C_*α*_ representation to run simulations of 56 IDPs and 14 MDPs (***Table S1, Table S2, Table S3***). In all systems, the interaction sites are located at the C_*α*_ positions in both folded and disordered regions; for the MDPs, we applied an additional elastic network model to keep domains intact during simulations (***Figure 1***A, see Methods). We term this combination of the force field parameters (CALVADOS 2) and the C_*α*_ representation of the interaction sites in the folded domains as CALVADOS2_C*α*_. As expected and reported previously (***Tesei and Lindorff-Larsen, 2023***), we found that simulations of IDPs with CALVADOS2_C*α*_ resulted in good agreement between experimental and calculated values of *R*_*g*_ (***Figure 1***B). In contrast, we found more substantial differences between experimental and calculated values of *R*_*g*_ for several MDPs (***Figure 1***B). In particular, we found that the *R*_*g*_ was underestimated for several MDPs including a series of two fluorescent proteins connected by Gly-Ser linkers of different lengths (here termed GS-proteins; ***Moses et al***. (***2024***)). This observation was confirmed by calculations of the relative mean signed deviation, rMSD, between experimental and calculated values of *R*_*g*_ that shows that these are on average underestimated by 18% in the MDPs (***Figure 1***B).

**Figure 1.**
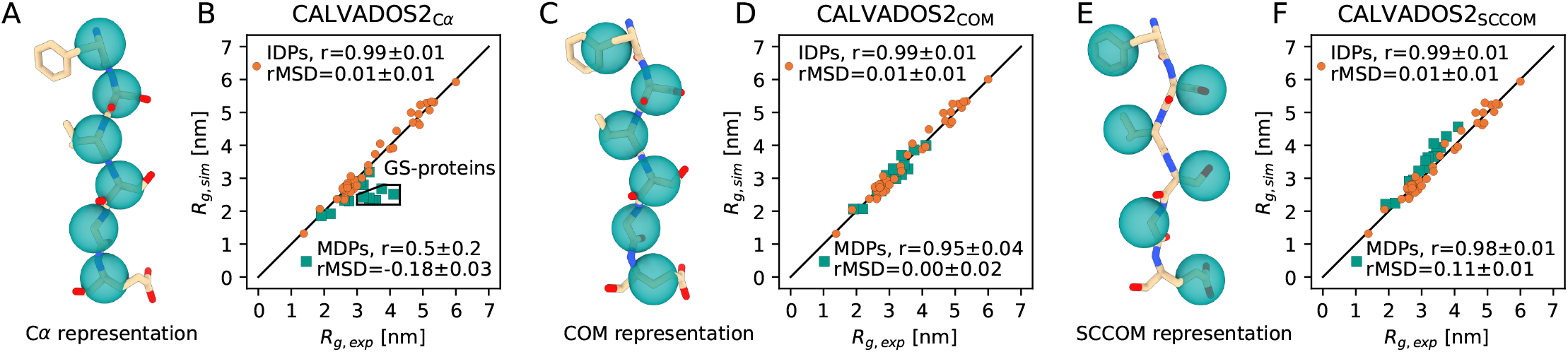
Simulations of MDPs and IDPs using a C_*α*_ representation, COM representation or side-chain centre-of-mass (SCCOM) representation. Location of the interaction sites in a *β*-sheet when using (A) a C_*α*_ representation, (C) a COM representation, and (E) a SCCOM representation. Comparison between simulated and experimental *R*_g_ values for IDPs (orange) and MDPs (green) using (B) the CALVADOS2_C*α*_ model (CALVADOS 2 parameters and a C_*α*_ representation for both folded and disordered regions), (D) the CALVADOS2_COM_ model (CALVADOS 2 parameters and a COM representation for the interaction sites in the folded regions), and (F) the CALVADOS2_SCCOM_ model (CALVADOS 2 parameters and a SCCOM representation for the interaction sites in the folded regions). The region labelled ‘GS-proteins’ in panel B contains a number of proteins consisting of pairs of */3*-sheet-rich fluorescent protein connected by glycine-serine linkers (***Moses et al., 2024***). Pearson correlation coefficients (*r*) and relative mean signed deviation rMSD = β(*R*_g,sim_ − *R*_g,exp_) / *R*_g,exp_β are reported in the legend, and errors represent standard errors of the mean calculated using bootstrapping. A negative rMSD value indicates that the calculated radii of gyration are systematically lower than the experimental values. The black diagonal lines in panel B, D and F indicate *y* = *x*.

As a first attempt at creating a model for both IDPs and MDPs, we used our previously described protocol (***Norgaard et al., 2008; Tesei et al., 2021b***) to optimize the *λ* stickiness parameters of the CALVADOS model targeting simultaneously SAXS and NMR data on 56 IDPs and 14 MDPs. The resulting *λ* values were generally smaller than those in CALVADOS 2 (***Figure S1***A) in line with the finding that the MDPs were too compact using CALVADOS 2. Nevertheless, it was also clear that this new parameter set made the agreement worse for disordered proteins (***Figure S1***B-E) and did not result in a satisfactory model to describe both IDPs and MDPs.

We instead hypothesized that the compaction of several MDPs was a result of placing the interaction sites at the C _*α*_ positions in the folded domains. In particular for *β* -sheet-containing proteins, this geometry would mean that residues whose side chains are buried inside the folded domain are represented by interaction sites located closer to the protein surface (***Figure 1***A); thus buried hydrophobic residues might appear as solvent exposed. We therefore constructed a new model where the interaction sites within folded regions were placed at the COM of the residue (***Figure 1***C) and constrained by harmonic restraints; when used with the CALVADOS 2 parameters, we term this model CALVADOS2_COM_. We stress that only the bead locations in the folded domains differ between the CALVADOS2_C*α*_ and CALVADOS2_COM_ models; residues in disordered regions are represented by one bead centred on the C_*α*_ positions in both models. In the absence of folded domains, CALVADOS2_COM_ and CALVADOS2_C*α*_ are thus identical and simulations with the two models gave comparable results (***Figure 1***B and D). In contrast, simulations of the MDPs with CALVADOS2_COM_ were in substantially better agreement with experiments than simulations with CALVADOS2_C*α*_ as evidenced e.g. by an increase in Pearson correlation coeicient from 0.5 to 0.95 and an increase in rMSD from -18% to 0% (***Figure 1***B and D). In addition to the COM representation, we also examined whether a side-chain centre-of-mass (SCCOM) representation, shifting bead positions of buried residues further away from the surface, could yield even more accurate *R*_*g*_ predictions than the COM representation (***Figure 1***E). We performed single chain simulations with the CALVADOS 2 parameters and the SCCOM representation (CALVADOS2_SCCOM_) and found that CALVADOS2_SCCOM_ on average resulted in an overestimation of the *R*_*g*_ of MDPs of 11% (***Figure 1***D and F). As an alternative solution to decrease the too strong interactions between folded domains, it has previously been suggested to scale down interactions between pairs of folded domains (by a factor of 0.7) and between folded domains and disordered regions (by a factor of 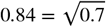) (***Krainer et al., 2021***). While applying this rescaling to CALVADOS 2 (termed CALVADOS2_C*α*_ 70%) led to improved agreement with experiments, the improvement was smaller than when using the COM representation, and the simulations had a remaining bias towards underestimating the radii of gyration (***Figure S2***). Therefore, we proceeded by using the COM representation in this study.

To examine in more detail why the CALVADOS2_C*α*_ model resulted in more compact conformations of MDPs than CALVADOS2_COM_, we calculated the time-averaged non-ionic (Ashbaugh-Hatch) interaction energies between residues of different folded domains. For this analysis we selected GS0, a construct with two fluorescent proteins separated by a 29-residue-long linker (***Moses et al., 2024***), since the *R*_*g*_ value of GS0 deviates substantially from experiments in simulations with CALVADOS2_C*α*_ (***Figure 1***B). In the energy maps, we see evidence of substantial inter-domain interactions between residue 140–230 of one fluorescent protein and residue 340– 440 of the other (***Figure 2***A). In contrast, these domain-domain interactions are not observed when simulating with COM representation (***Figure 2***B). The comparison of the two energy maps thus supports the hypothesis that the too compact conformations of MDPs in simulations with CALVADOS2_C*α*_ result from inter-domain attractions that are decreased in the COM representation (***Figure 2***C).

**Figure 2.**
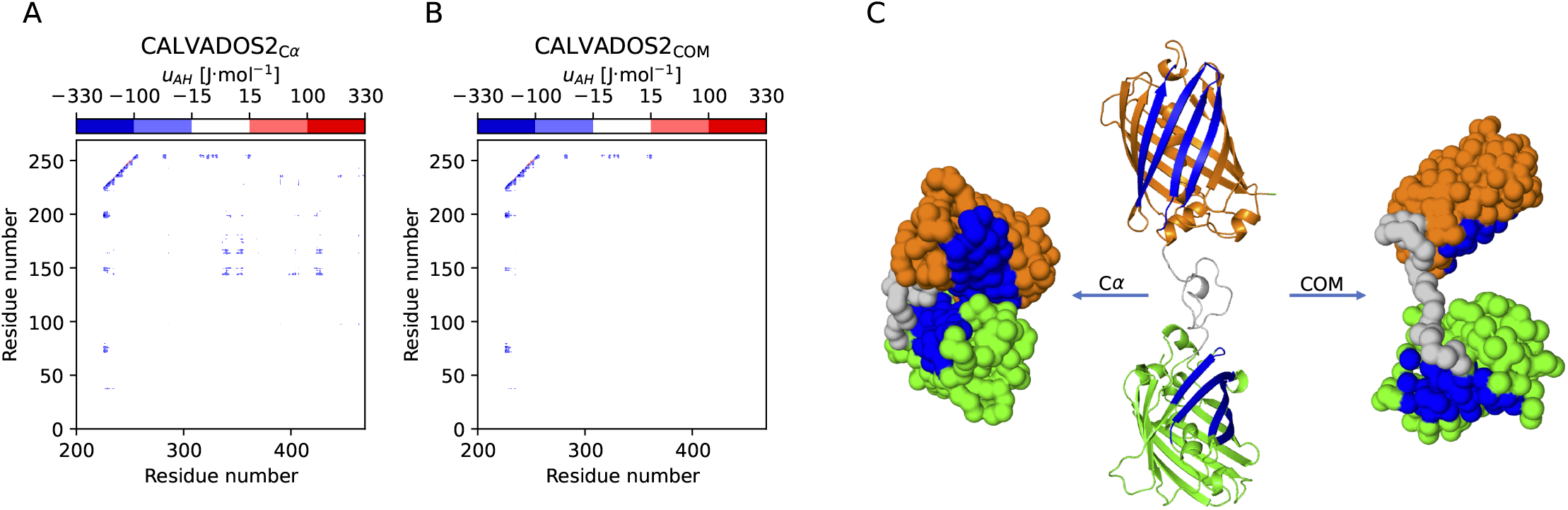
Energy calculations reveal substantial inter-domain interactions. We calculated interaction energy maps (of the Ashbaugh-Hatch term in the force field) from simulations using (A) the CALVADOS2_C*α*_ model and (B) the CALVADOS2_COM_ model. We show only a subset of the map representing interactions between the first (residues 1–226 on the y-axis) and second (residues 256–470 on the x-axis) folded domains. (C) Examples of structures of GS0 with the same *R*_*g*_ as the average over simulations using CALVADOS2_C*α*_ (left) and CALVADOS2_COM_ (right). The starting structure of the simulations is shown in the middle, where green and orange parts are the two fluorescent proteins connected by a flexible linker (grey). The regions that interact strongly in the CALVADOS2_C*α*_ simulations are coloured blue.

### Optimizing CALVADOS using a centre-of-mass representation

Having shown that the COM representation gave an improved description of MDPs while preserving the accuracy when simulating IDPs, we proceeded to optimize the CALVADOS model further. We used our iterative Bayesian optimization scheme (***Norgaard et al., 2008; Tesei et al., 2021b***) to optimize the *λ* stickiness parameters of the CALVADOS model targeting simultaneously SAXS and NMR data on 56 IDPs and 14 MDPs (***Table S1, Table S2, Table S3***). In these simulations we used the COM representation of the folded domains and we thus term the final model CALVADOS3_COM_ to represent both the force field and the COM representation of the folded regions. The resulting *λ* values in CALVADOS3_COM_ are similar to those in CALVADOS 2 (***Figure 3*** and ***Figure S3***). We found that simulations of IDPs with CALVADOS3_COM_ and CALVADOS2_COM_ gave similar agreement to SAXS experiments. Likewise, we found a similar agreement for the MDPs (***Figure 1***D, ***Figure 3***B and C).

**Figure 3.**
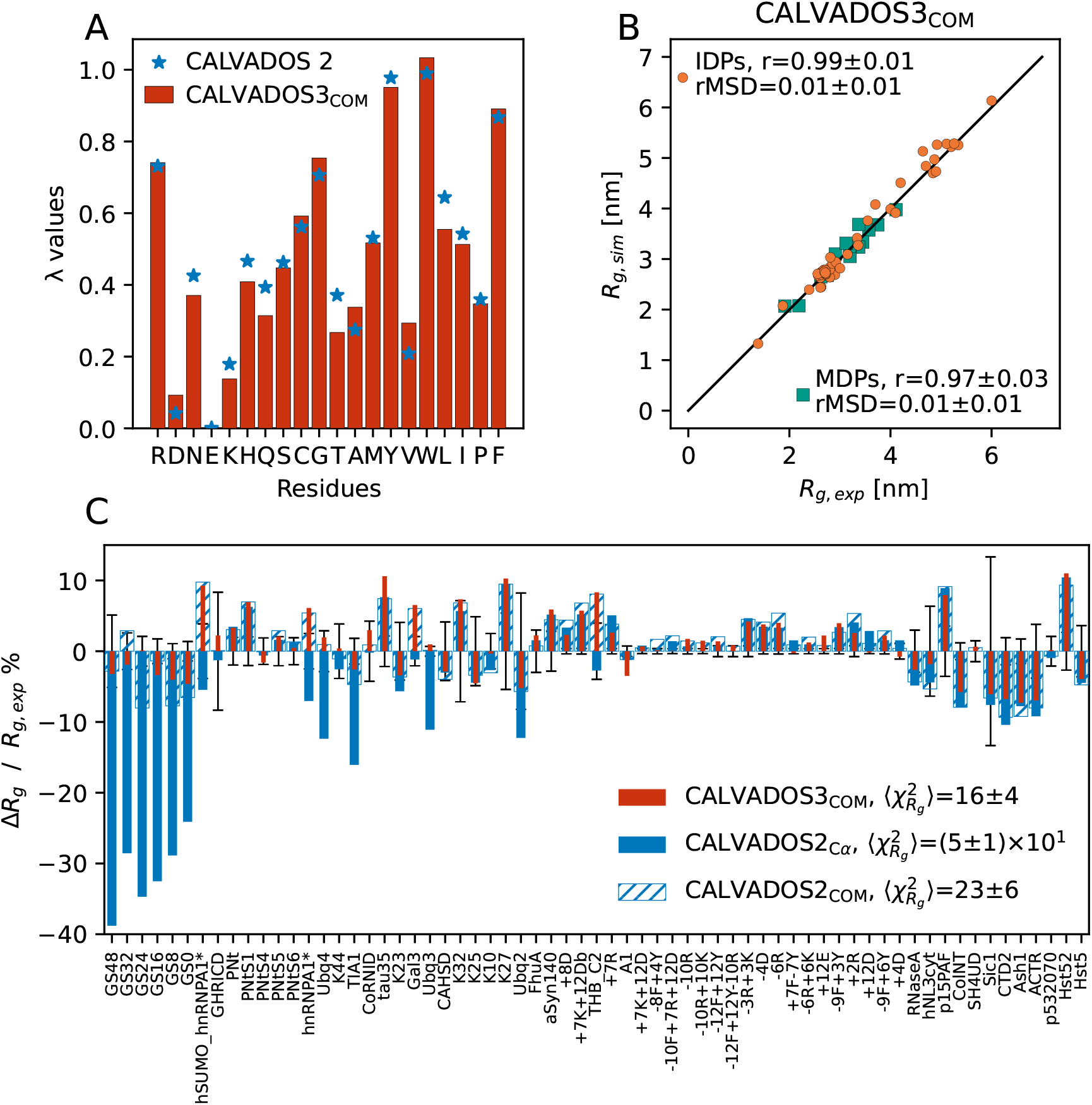
Optimizing the *λ* parameters using a COM representation for folded domains. (A) Comparison between *λ* values from CALVADOS 2 (blue) and CALVADOS3_COM_ (red). (B) Comparison between simulated and experimental *R*_*g*_ values for IDPs (orange) and MDPs (green) using CALVADOS3_COM_. Pearson correlation coeicients (*r*) and rMSD are reported in the legend. The black diagonal line indicates *y* = *x*. (C) Relative difference between experimental and simulated *R*_*g*_ values from CALVADOS3_COM_ (red), CALVADOS2_C*α*_ (blue) and CALVADOS2_COM_ (blue hatched). 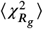 values across IDPs and MDPs in training set are reported in the legend. Error bars show the experimental error divided by *R*_g,exp_.

Having optimized *λ*, we validated the CALVADOS3_COM_ model on 25 IDPs and 9 MDPs (***Table S4, Table S5***) that were not used in training for any of the models (***Figure 4***). For the 25 IDPs we found good agreement for all three models (CALVADOS2_C*α*_, CALVADOS2_COM_ and CALVADOS3_COM_) (***Figure 4***A–C). We note again that the COM representation is only applied to the folded domain. All IDPs have C_*α*_ representations, so CALVADOS2_C*α*_ and CALVADOS2_COM_ are the same models for IDPs. In contrast, for MDPs we found that CALVADOS3_COM_ and CALVADOS2_COM_ perform substantially better than CALVADOS2_C*α*_ (***Figure 4***A-C). Our validation results thus show that the CALVADOS3_COM_ model gives improved agreement for simulations of MDPs while retaining the accuracy of CALVADOS2_C*α*_ for simulations of IDPs. Across the 34 independent test proteins we find 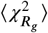 values of 50, 22 and 15 for CALVADOS2_C*α*_, CALVADOS2_COM_ and CALVADOS3_COM_, respectively (***Figure S4***), and both CALVADOS2_COM_ and CALVADOS3_COM_ have essentially no bias (rMSD≈0; ***Figure 4***B and C).

**Figure 4.**
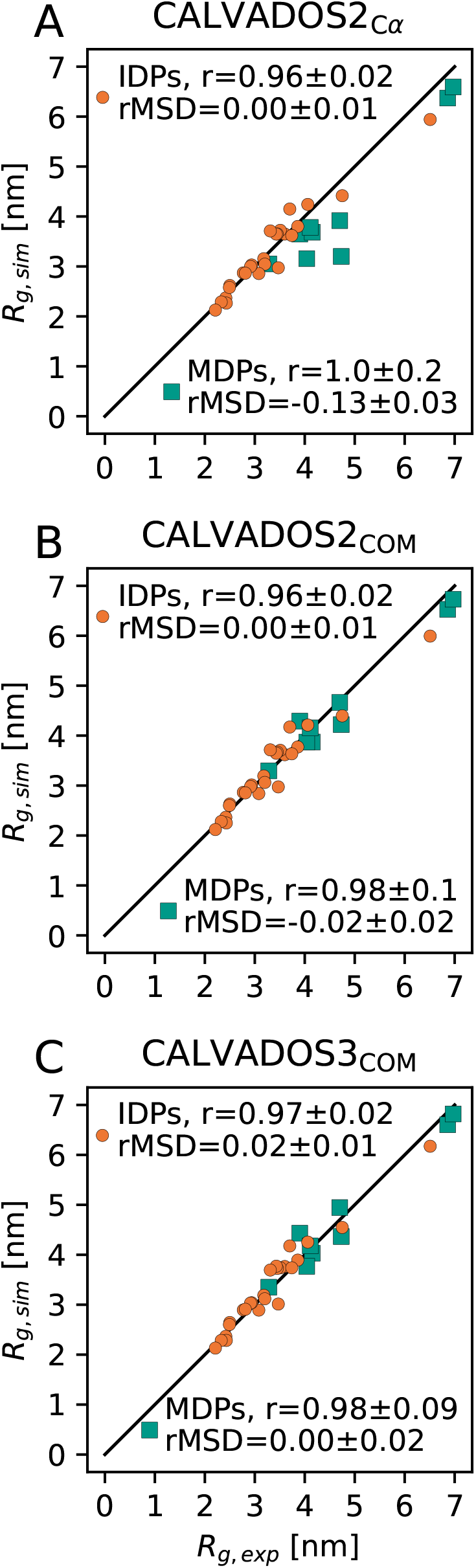
Validation of the CALVADOS3_COM_ model using proteins that were not used during training. Comparison of simulated and experimental *R*_*g*_ values on a validation set using (A) CALVADOS2_C*α*_, (B) CALVADOS2_COM_ and (C) CALVADOS3_COM_. Pearson correlation coeicients (*r*) and rMSD are reported in the legend. The black diagonal lines indicate *y* = *x*.

### Simulations of phase separation of disordered and multi-domain proteins

We and others have previously used one-bead-per-residue models such as CALVADOS to study the self-association and phase separation of IDPs (***Dignon et al., 2018; Tesei et al., 2021b; Joseph et al., 2021; Regy et al., 2021; Dannenhoffer-Lafage and Best, 2021; Wessén et al., 2022; Tesei and Lindorff-Larsen, 2023; Valdes-Garcia et al., 2023***). In some cases, these models have also been used to study phase separation of proteins that contain a mixture of folded and disordered regions (***Dignon et al., 2018; Conicella et al., 2020; Her et al., 2022***). We therefore examined whether the CALVADOS3_COM_ model could be used to study phase separation of both IDPs and MDPs. We used multi-chain simulations in a slab geometry (***Dignon et al., 2018***) to simulate the partitioning of proteins between a dilute and dense phase, and calculated the dilute phase concentration (the saturation concentration; *c*_sat_) as a sensitive measure of the accuracy of the model. We first simulated 33 IDPs and found that simulations with CALVADOS3_COM_ gave an agreement with experimental values of *c*_sat_ that is comparable to that of CALVADOS2_C*α*_ (***Table S6, Figure S5, Figure S6, Figure S7***).

We then proceeded to use CALVADOS3_COM_ to study the phase separation of MDPs including hnRNPA1* (where * denotes that residues 259–264 have been deleted from full-length hnRNPA1), full-length FUS (FL_FUS) and other multi-domain proteins with experimental estimates of *c*_sat_ (***Table S7; Wang et al***. (***2018***); ***Martin et al***. (***2021b***)). Simulations of hnRNPA1* with CALVADOS2_C*α*_, under conditions where the experimental dilute phase concentration is 0.17 mM, resulted in essentially all proteins in the dense phase (*c*_sat_=0 mM; ***Figure 5***A). In contrast, simulations using CALVADOS3_COM_ resulted in a lower propensity to phase separate and a calculated value of *c*_sat_=0.14±0.01 mM that is comparable to experiments (***Figure 5***B).

**Figure 5.**
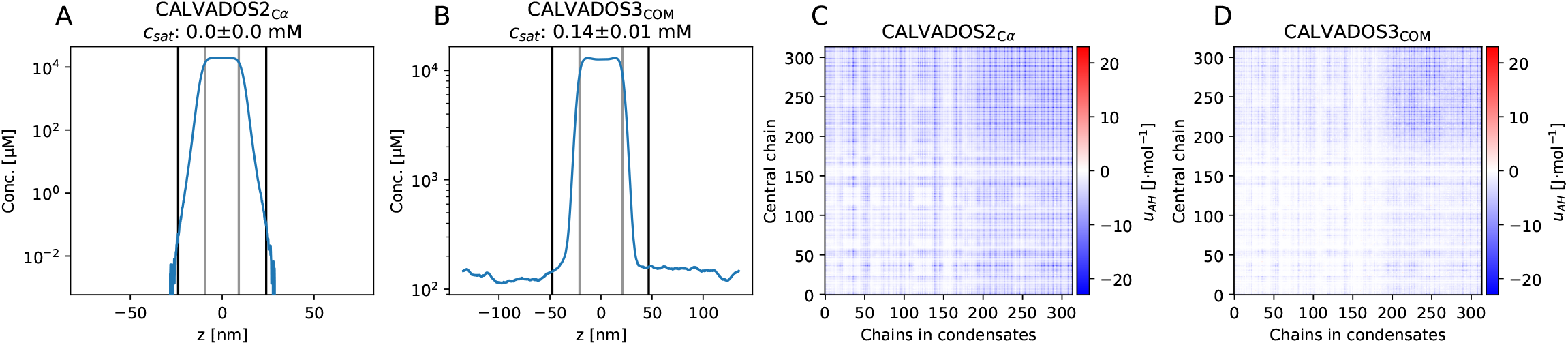
Phase coexistence simulations of hnRNPA1* using (A, C) CALVADOS2_C*α*_ and (B, D) CALVADOS3_COM_. Simulations were performed at 293 K and an ionic strength of 0.15 M. Equilibrium density profile of hnRNPA1* using (A) CALVADOS2_C*α*_ and (B) CALVADOS3_COM_. *c*_sat_ calculated from density profiles are 0 mM and 0.14 mM, respectively. Average residue-residue interaction energies (the Ashbaugh-Hatch term in the force field) between the most central chain and the rest of the condensate for (C) CALVADOS2_C*α*_ and (D) CALVADOS3_COM_.

To understand the origin of these differences we calculated interaction energy maps of the proteins in the dense phase. Experiments have shown that the LCD in hnRNPA1* (residues 186–320) plays a central role in driving phase separation (***Molliex et al., 2015; Martin et al., 2021b***), and we indeed found evidence for substantial LCD-LCD interactions in the dense phases in simulations with both CALVADOS2_C*α*_ (***Figure 5***C) and CALVADOS3_COM_ (***Figure 5***D). In the simulations with CALVADOS2_C*α*_ we, however, also observed more sub-stantial interactions between the folded RRM (RNA recognition motif) domains (residues 14–97 and 105–185) and between the RRMs and the LCD. In simulations with CALVADOS3_COM_ these interactions were much weaker, presumably explaining the increase of *c*_sat_ in these simulations.

Having demonstrated that CALVADOS3_COM_ provides a more accurate description of the phase behaviour of hnRNPA1* than CALVADOS2_C*α*_, we proceeded to perform simulations of several other MDPs for which we found estimates of *c*_sat_ in the literature (***Figure 6, Figure S8, Figure S9***). As for hnRNPA1*, we found that CALVADOS2_C*α*_ substantially overestimates the tendency of these proteins to undergo phase separation (i.e. underestimate *c*_sat_). The use of the COM representation in CALVADOS3_COM_ decreases the protein-protein interactions, and thus substantially improves the agreement with experiments, though differences remain.

**Figure 6.**
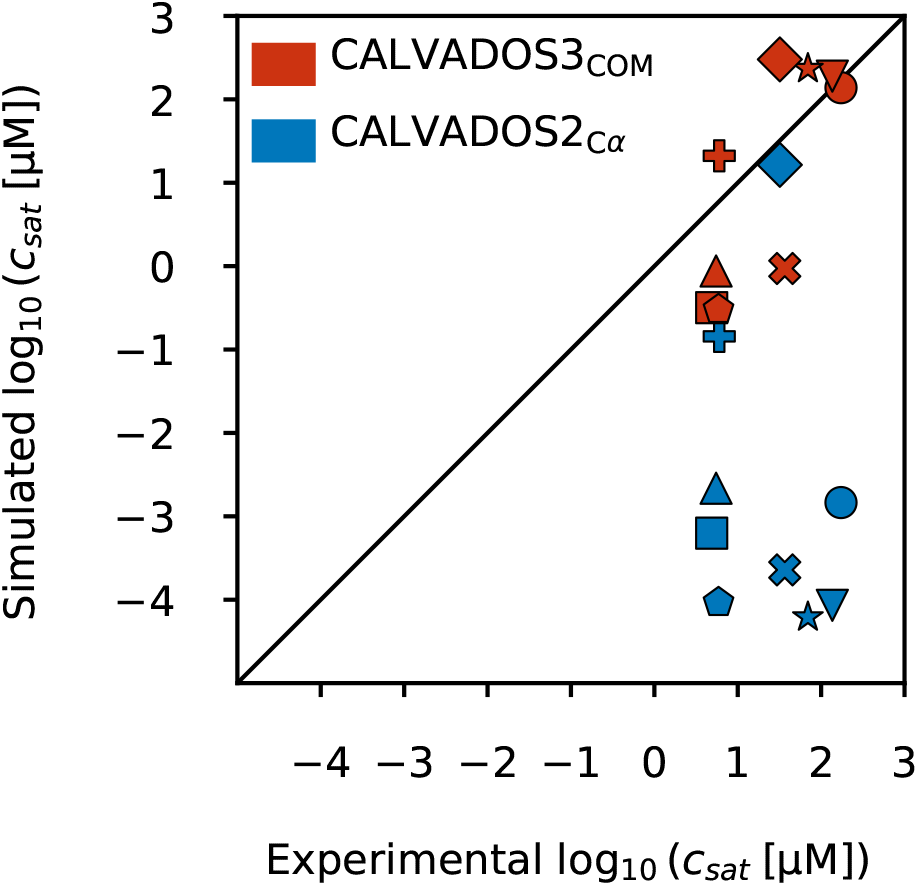
Comparison between simulated and experimental *c*_sat_ values for MDPs using the CALVADOS3_COM_ model (red) and CALVADOS2_C*α*_ (blue). The simulated proteins are hnRNPA1* (circle), hSUMO_hnRNPA1* (downward triangle), FL_FUS (upward triangle), GFP_FUS (square), SNAP_FUS (pentagon), SNAP_FUS_PLDY2F_RBDR2K (star), SNAP_FUS_PLDY2F (x symbol), FUS_PLDY2F_RBDR2K (diamond) and hnRNPA3 (plus symbol). The black diagonal line indicates *y* = *x*.

### Examining changes in folding stability in condensates

Experiments have shown that the protein-rich environment of condensates can modulate the stability of folded proteins or nucleic acids (***Nott et al., 2015; Ruff et al., 2022; Chen et al., 2024; Ahmed et al., 2024***). Inspired by these findings, we used the ability to simulate both folded and disordered regions with CALVADOS 3 to examine how partitioning into condensates may shift the folding equilibrium of a folded domain. As it is diicult to sample the folding-unfolding equilibrium by simulations, we studied it indirectly using a thermodynamic cycle that involves differences in partitioning of the folded and unfolded forms into a condensate (***Nott et al., 2015***).

To demonstrate how CALVADOS 3 enables such analyses, we simulated the isolated RRM1 and RRM2 from hnRNPA1^*^ (***Figure 7***A) in the presence of a condensate of the LCD of hnRNPA1^*^ and calculated the free energies of partitioning of the RRM domains in their native, folded state, 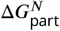. Using the same approach, we performed direct-coexistence simulations without applying harmonic networks to the RRMs to calculate the free energies of partitioning of the RMMs in their unfolded state, 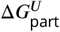. A comparison of the concentration profiles from our direct-coexistence simulations shows that the unfolded states accumulate in the condensate and are depleted from the dilute phase to a greater extent than the folded states (***Figure 7***B–C); We quantify this via a more negative free energy of partitioning, 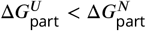 (***Figure 7***D). The preference of the unfolded state for the condensate is particularly pronounced for RRM2, for which we estimate a two-fold decrease in the free energy of partitioning 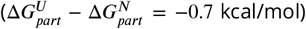. From the thermodynamic cycle, this in turn means that the folding stability of RRM2 is 0.7 kcal/mol lower (less stable) in the condensate than in the dilute phase.

**Figure 7.**
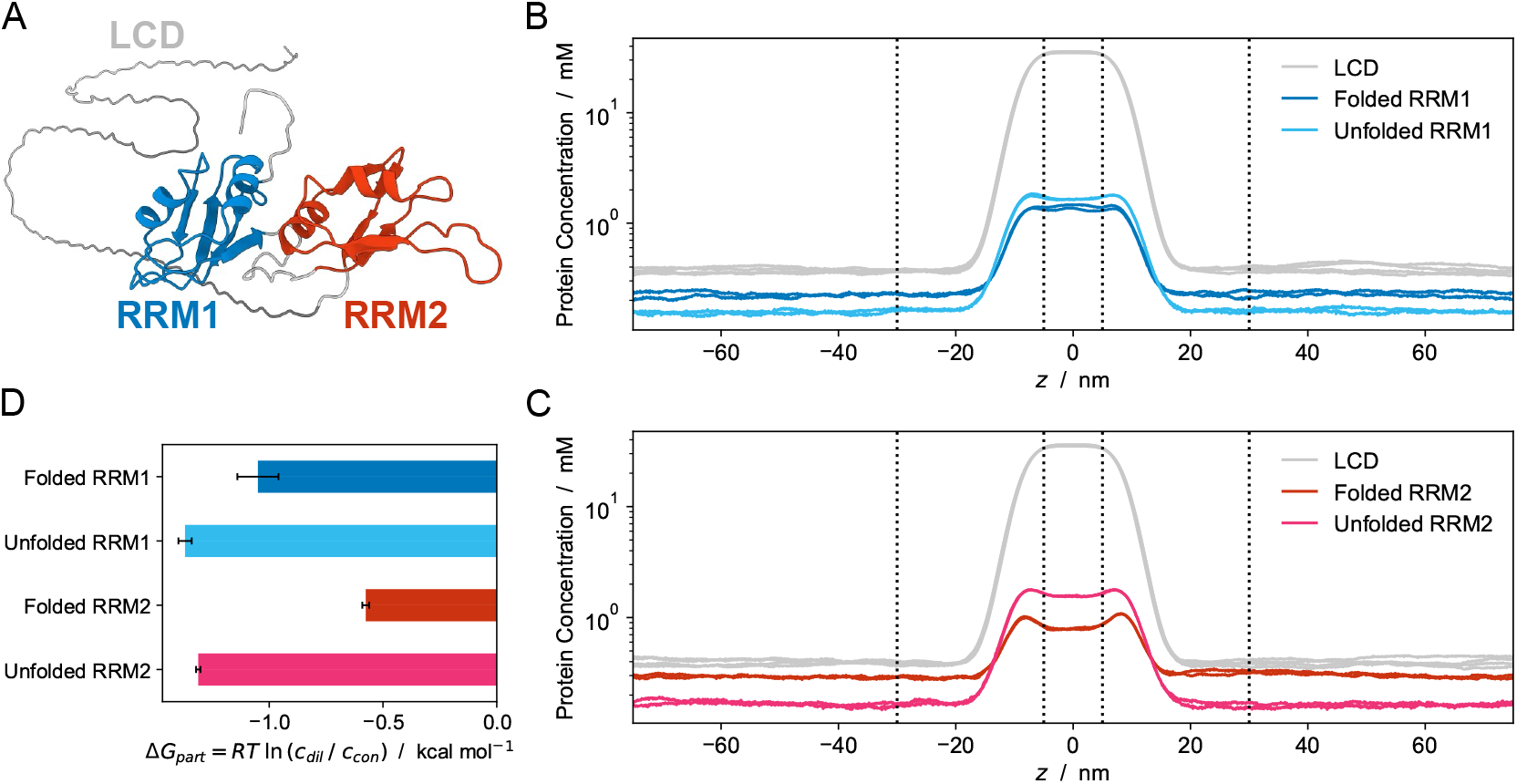
Predicting the effect of the protein-rich environment of a condensate on the stability of folded domains. (A) Structure of hnRNPA1^*^ highlighting the low-complexity domain (grey) and RNA-recognition motifs 1 (blue) and 2 (red). (B) Concentration profiles of the LCD (grey) and RRM1 in the native (blue) and unfolded (cyan) state. (C) Concentration profiles of the LCD (grey) and RRM2 in the native (red) and unfolded (magenta) state. (D) Free energy of partitioning of RRM1 and RRM2 in native and unfolded states into condensates of the LCD. Data estimated from direct-coexistence simulations performed in two independent replicates. Error bars in (D) represent the differences between the replicates.

To put these changes into context, we used a recently developed machine learning approach (***Cagiada et al., 2024***) to predict the absolute protein folding stabilities of the isolated RRMs in the dilute phase, 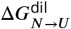, and obtained 6.6 kcal/mol for RRM1 and 4.4 kcal/mol for RRM2. Using these values and assuming a two-state model, we estimate that the partitioning into the condensate has a negligible effect on the amount of unfolded state for RRM1; in contrast we predict a four-fold increase in the population of the unfolded state of RRM2 from 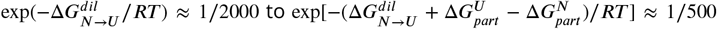. Although substantial additional work is needed to examine the accuracy of CALVADOS 3 for quantifying differences in partitioning of folded and unfolded proteins into condensates, these data show a promising use of our model for predicting unfolding in condensates.

## Discussion

In this work, we found that simulations with the CALVADOS2_C*α*_ model, previously shown to represent single-chain and multi-chain properties of IDPs, underestimated the radii of gyration of MDPs. Changing the CG mapping method from C_*α*_ to COM substantially improved the agreement with experimental data. This observation is in line with the finding that reconstruction of all-atom structures from a centre-of-mass representation is more accurate than from a C_*α*_ representation (***Heo and Feig***, ***2024***). We reoptimized the ‘stickiness’ parameters in the context of a COM-based model based on experimental data for both IDPs and MDPs. The resulting CALVADOS3_COM_ model provides a good description of both single- and multi-chain simulations of both IDPs and MDPs.

The relatively low *c*_sat_ value calculated from slab simulations of hnRNPA1* with CALVADOS2_C*α*_ further supported that interactions between the folded domains are overestimated by C_*α*_ -based models without any further modifications. Considering that the SCCOM-based model (CALVADOS2_SCCOM_) overestimated *R*_*g*_ of MDPs, we suggest that the COM-based model (CALVADOS3_COM_) appears to strike a good balance, leading to improved values of *c*_sat_ for MDPs. Nevertheless, some systematic differences remain even with this model, which resulted in underestimates of *c*_sat_ for different constructs of the protein FUS. Together, our results show that the new parameter set and the centre-of-mass representation (CALVADOS3_COM_) retain the accuracy of CALVADOS 2 for IDPs, but improve the description of proteins with both disordered and folded domains. We therefore term this new model CALVADOS 3, with the implicit notion that this model is used with centre-of-mass representation of residues within folded regions. We note that an earlier version of this preprint (***Cao et al., 2024***) used a slightly different set of parameters, and we suggest to refer to that model as CALVADOS 3beta.

When simulating MDPs with CALVADOS 3 we need to restrain the folded domains using harmonic restraints. In the current work we have manually determined the boundaries for which regions are considered to be folded, though automated methods will be needed for large-scale applications. Tools for automatic predictions of domain boundaries exist (***Holm and Sander***, ***1994; Lau et al., 2023***) and might be combined with AlphaFold to set the harmonic restraints (***Jussupow and Kaila***, ***2023***).

Despite these current limitations, we envision that the CALVADOS 3 model will enable detailed studies of the interactions within and between multi-domain proteins, and pave the way for proteome-wide simulation studies of full-length proteins similar to what has recently been achieved for IDRs (***Tesei et al., 2024***). We also envision that our approach to study changes in protein stability inside condensates can be used together with methods to predict absolute protein stability (***Cagiada et al., 2024***) to learn and expand our knowledge on the rules that underlie phase separation and changes in stability of folded, globular proteins (***Ruff et al., 2022***).

## Methods

### Description of the model

We modelled each amino acid by one bead. We generated C_*α*_ -beads for IDPs and assigned C_*α*_ atom coordinates to bead positions for IDRs in multi-domain proteins according to their modelled or experimental structures (see below, Simulations). For structured domains, we used the following rules for the different representations: we placed each bead position at the C_*α*_ atom (C_*α*_ representation), or the centre of mass calculated for all the atoms in a residue (COM representation), or the centre of mass calculated for only side chain atoms of a residue (SC-COM representation). The CALVADOS 3 energy function consists of bonded interactions, non-bonded interactions and an elastic network model as described below.

Chain connectivity of the beads is described by a harmonic potential,

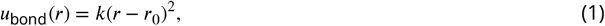

with force constant *k* = 8033 kJ·mol^−1^-nm^−2^. The equilibrium distance *r*_0_ is set to 0.38 nm if two beads are both within IDRs, or the distance between two beads in the initial conformation if at least one bead is within a folded domain.

For non-bonded interactions, we use a truncated and shifted Ashbaugh-Hatch (AH) and Debye-Hückel (DH) potential to model van der Waals and salt-screened electrostatic interactions, respectively. The Ashbaugh-Hatch potential is described by

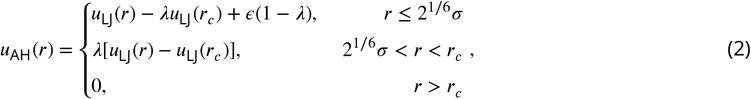

where *u*_LJ_(*r*) is the Lennard-Jones (LJ) potential,

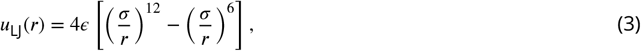

and where *ϵ* = 0.8368 kJ·mol^−1^ and *r*_*c*_ = 2.2 or 2 nm. Similar to previous work, we use *r*_*c*_ = 2.2 nm during the optimization of CALVADOS3_COM_, and use 2 nm during validation and application (***Tesei and Lindorff-Larsen***, ***2023***). Both *σ* and *λ* are calculated as the arithmetic averages of residue-specific bead size and stickiness, respectively. *σ* values are van der Waals volumes calculated by Kim and Hummer (***Kim and Hummer***, ***2008***). *λ* values are treated as free parameters and optimized iteratively through a Bayesian parameter-learning procedure as described previously (***Tesei et al., 2021b; Tesei and Lindorff-Larsen***, ***2023***) to minimize the differences in the simulated and experimental *R*_*g*_ and PRE data. In simulations where we scaled down interactions of folded domains (CALVADOS2_C*α*_ 70%)), we scaled down *ϵ* to 0.7*ϵ* for domain-domain interactions and to 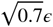 for domain-IDR interactions.

The Debye-Hückel potential is described by

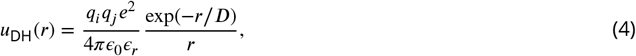

where *q* is the average amino acid charge number, *e* is the elementary charge, 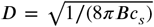 is the Debye length of an electrolyte solution of ionic strength *c*_*s*_, *B*(*ϵ*_*r*_) is the Bjerrum length and *ϵ*_0_ is vacuum permittivity. Electrostatic interactions are truncated and shifted at the cutoff distance *r*_*c*_ = 4 nm. The temperature-dependent dielectric constant of the implicit aqueous solution is modelled by the following empirical relationship (***Akerlof and Oshry***, ***1950***):

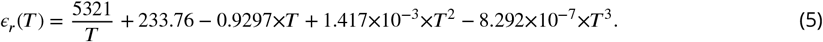

we use the Henderson-Hasselbalch equation to estimate the average charge of the histidine residues, assuming a *p****K***_*a*_ value of 6 (***Nagai et al., 2008***).

We use an elastic network model (ENM) with a harmonic potential to restrain non-bonded pairs in the folded domains using

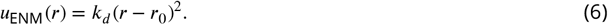

Here, the force constant *k*_*d*_ is 700 kJ·mol^−1^-nm^−2^, *r* is the distance between beads and equilibrium distances *r*_0_ are directly taken from the reference structures. We only apply the ENM to residue pairs with an *r*_0_ below a 0.9 nm cutoff. We determine the predefined boundary of each domain in MDPs by visual inspection of the three-dimensional structures (***Table S8***). Each domain has a starting amino acid and an ending amino acid indicating the range of the domain. Only residue pairs within the same domain are restrained by this harmonic potential except for bonded pairs, which are restrained by the aforementioned bonded potential. All boundaries of MDPs are consistent with definitions in their experimental or simulation articles. In some cases, one domain could be discontinuous because of long loops within the domain so we exclude those regions when defining boundaries. Residues of *α* -helix, *β*-sheet and short loops in a structured domain are all restrained equally with the same force constant and cutoff distance. The application of ENM ensures that secondary structures within folded domains would not fluctuate substantially (***Figure S10***). Non-bonded interactions (Ashbaugh-Hatch and Debye-Hückel potential) are excluded for the restrained pairs.

### Simulations

We generated initial conformations of all IDPs as Archimedes’ spirals with a distance of 0.38 nm between bonded beads. Atomistic structures of all MDPs used in optimization procedures, single-chain validation and slab simulations either came from our recent work (***Thomasen et al., 2023***) or were modelled by superposing experimental domain structures (if available) on AlphaFold predictions (***Jumper et al., 2021; Varadi et al., 2022***). We then mapped all of these MDPs to CG structures based on different CG representations (C_*α*_, COM, SCCOM).

We conducted Langevin dynamics simulations using OpenMM 7.6.0 (***Eastman et al., 2017***) in the NVT ensemble with an integration time step of 10 fs and friction coefficient of 0.01 ps^−1^. Single chains of *N* residues were simulated in a cubic box with a (*N* − 1)×0.38 + 4 nm box edge length under periodic boundary conditions. Each chain was simulated in 20 replicas for 6.3∼77.7 ns depending on the sequence length of the disordered regions (***Tesei and Lindorff-Larsen***, ***2023; Tesei et al., 2024***). Final trajectories had 4000 frames for each protein, excluding the initial 10 frames in each replica.

We performed direct-coexistence simulations in a cuboidal box using [*L*_*x*_, *L*_*y*_, *L*_*z*_] = [17, 17, 300] and [15, 15, 150] nm to simulate multi-chains of Ddx4WT and the other IDPs, respectively. For MDPs, box sizes are shown in ***Table S7***. To keep the condensates thick enough and reduce finite-size surface effects, we chose 150 chains for hnRNPA1* and 100 chains for all the other IDPs and MDPs (see also below). We generated each IDP chain as an Archimedes spiral with a distance of 0.38 nm between bonded beads in the xy-plane. Each spiral was placed along the z-axis with a 1.47 nm interval. To avoid steric clashes of densely packed MDP input structures, we chose the most compact conformation sampled by single-chain simulations with CALVADOS 2 parameters and corresponding CG representation as the initial conformation for each MDP chain. Before production simulations, we performed equilibrium runs where we used an external force to push each chain towards the centre of the box so that a condensate could be formed. We then continued to perform production simulations, saving frames every 0.125 ns and discarded the first 150 ns before analysis. The slab in each frame was centred in the box and the equilibrium density profile *ρ* (*z*) was calculated by taking the averaged densities over the trajectories as previously described (***Tesei and Lindorff-Larsen***, ***2023***).

To examine finite-size effects of the direct-coexistence simulations we performed additional simulations of hnRNPA1* varying both the box dimensions (*L*_*x*_, *L*_*y*_, *L*_*z*_) and the number of chains. We calculated both dense and dilute phase concentrations from each simulation and find that unless we use a very small patch (*L*_*x*_ = *L*_*y*_ = 11 nm), the results are consistent (***Figure S11, Figure S12, Table S9***), in line with previous analyses of such finite-size effects (***Dignon et al., 2018; Joseph et al., 2021***). Convergence of the IDP simulations was assessed as previously described (***Tesei et al., 2021b***).

To indicate the computational performance of single- and multi-chain CALVADOS simulations, we show the performance for systems of different sizes run either on an Intel Xeon Gold 6130 CPU (for single-chain simulations) on an NVIDIA Tesla V100 GPU (for multi-chain simulations) (***Figure S13***).

To estimate the free energy of partitioning of RRM1 (residues 11–89) and RRM2 (residues 105–179) into condensates of hnRNPA1^*^ LCD (GS followed by residues 186–314), we performed direct-coexistence simulations at 298 K, pH 7.5, and 150 mM ionic strength, in a cuboidal box with sidelengths [*L*_*x*_, *L*_*y*_, *L*_*z*_] = [15, 15, 150] nm. The structures of the native states of RRM1 and RRM2 were based on the crystal structure (***Shamoo et al., 1997***) as previously described (***Martin et al., 2021b***). We performed two independent simulations, each 21 µs long, for each system and, after centering the LCD condensate in the middle of the box, calculated concentration profiles along the *z*-axis using the last 20 µs of each trajectory. We estimated the free energies of partitioning as Δ*G*_part_ = *RT* ln (*c*_dil_ / *c*_con_) where *R* is the gas constant and *c*_dil_ and *c*_con_ are the average concentrations of the RRMs in the dilute phase and in the LCD condensate, respectively. The error on Δ*G*_part_ was estimated as the difference between the values from the two independent simulation replicas. Absolute folding stabilities of RRM1 and RRM2 were calculated using the Google Colab implementation of a recently described model for predicting absolute protein stability (***Cagiada et al., 2024***).

### Parameter optimization

Our Bayesian Parameter-Learning Procedure (***Tesei and Lindorff-Larsen***, ***2023***) of the ‘stickiness’ parameters, *λ*, aimed to minimize the following cost function:

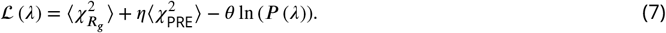

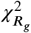 and 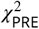 denoting *R*_*g*_ and PRE differences between experiments and simulations are estimated as

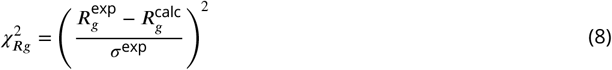

and

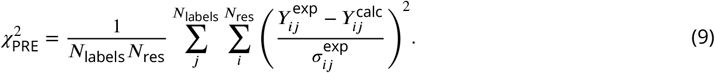

Here *P* (*λ*) is a statistical prior of *λ* (***Tesei et al., 2021b; Tesei and Lindorff-Larsen***, ***2023***), *(σ*^exp^ is the error on the experimental values, *Y* is PRE data, either *I*_para_/*I*_dia_ or r_2_ is calculated using the rotamer library approach implemented in DEER-PREdict (***Tesei et al., 2021a***), *N*_labels_ is the number of spin-labeled mutants, and *N*_*res*_ is the number of measured residues. The prior loss, θ ln (*P* (*λ*)), quantifies the difference between prior distribution *P* (*λ*) and current *λ* values (with min-max normalization at each step) to avoid overfitting. The coeicients are set to *η*, = 0.1 and *θ* = 0.08. *λ* is not allowed to be negative but can be greater than 1.0 during optimization.

We used a training set consisting of 56 IDPs and 14 MDPs to perform the optimization. All of those proteins were from our previous studies (***Tesei and Lindorff-Larsen***, ***2023; Thomasen et al., 2023***). A summary of the training data and other properties of different CALVADOS models is shown in the supporting material (***Table S10***). 51 IDPs and 14 MDPs in this training set were used for fitting against experimental SAXS *R*_*g*_ data and 5 IDPs were used for fitting against experimental PRE data (***Table S1, Table S2, Table S3***). We then used a validation set to validate the performances of our new optimized models on reproducing experimental *R*_*g*_. This validation set was composed of 25 IDPs and 9 MDPs. 12 IDPs in this validation set were from our previous work and the rest (13 IDPs and 9 MDPs) were newly collected experimental *R*_*g*_ data in this work (***Table S4, Table S5***). We also collected nine MDPs with measured values of *c*_sat_ to examine the accuracy of the phase behaviour simulated with the models presented in this work (***Table S7***).

The optimization procedure went through several cycles until convergence of the final total cost (| Δℒ| < 1, Δℒ is the difference of final total cost between the current and previous cycle, ***Equation 7***). Within each cycle, we use the optimized *λ* values from the previous cycle to perform new single-chain simulations (initial *λ* values for the first cycle are CALVADOS 2 parameters, (***Tesei and Lindorff-Larsen***, ***2023***)), calculate *R*_*g*_ and PRE for each frame and then nudge values in the *λ* set iteratively to minimize the cost function (five residues are randomly subjected to small perturbations sampled from a Gaussian distribution with *µ* = 0, *(σ* = 0.05). This trial *λ* set (*λ*_*k*_) is used to calculate the Boltzmann weights of each frame by *w*_*i*_ = exp(−[*U* (*r*_*i*_, *λ*_*k*_) − *U* (*r*_*i*_, *λ*_0_)]/*k*_B_*T*), where *U* is the AH potential, *r*_*i*_ are coordinates of a conformation, *k*_B_ is the Boltzmann constant and *T* is temperature. The resulting weights are then used to calculate the effective fraction of frames by 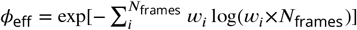; if *ϕ*_eff_ ≥ 0.6, trial *λ*_*k*_ acceptance probability is determined by the Metropolis criterion, 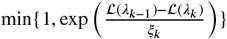, where *ξ*_*k*_ is a unitless control parameter, its initial value is set to 0.1 and scaled down by 1% at each iteration until *ξ <* 10^−8^, which means a micro-cycle is complete. Within a cycle, a total of 10 micro-cycle are performed. In this work, the optimization procedure converged within three cycles. Therefore, we used the resulting *λ* values from the third cycle as the final parameter set. We ran one additional optimization cycle to confirm the convergence of the training.

## Data and software availability

Scripts and data to reproduce the work are available via https://github.com/KULL-Centre/_2024_Cao_CALVADOSCOM.

## Acknowledgments

We acknowledge the use of computational resources from Computerome 2.0, the ROBUST Resource for Biomolecular Simulations (supported by the Novo Nordisk Foundation; NNF18OC0032608) and the Biocomputing Core Facility at the Department of Biology, University of Copenhagen. This research was supported by the PRISM (Protein Interactions and Stability in Medicine and Genomics) centre funded by the Novo Nordisk Foundation (NNF18OC0033950, to K.L.-L.), and CSC (China scholarship council, 202206340019). SB is a recipient of an EMBO postdoctoral fellowship (ALTF 810-2022).

---

**Table S1.** Experimental solution conditions and radii of gyration of IDPs included in the training set for the Bayesian parameter-learning procedure.

| Protein | N | $R_g \pm \text{Err}$ [nm] | T [K] | $c_s$ [M] | pH | Ref. |
| --- | --- | --- | --- | --- | --- | --- |
| Hst5 | 24 | $1.38 \pm 0.05$ | 293.00 | 0.150 | 7.5 | (Jephthah et al., 2019) |
| Hst52 | 48 | $1.87 \pm 0.05$ | 298.00 | 0.150 | 7.0 | (Fagerberg et al., 2020) |
| p532070 | 62 | $2.39 \pm 0.05$ | 277.00 | 0.100 | 7.0 | (Zhao et al., 2021) |
| ACTR | 71 | $2.63 \pm 0.1$ | 278.00 | 0.200 | 7.4 | (Kjaergaard et al., 2010) |
| Ash1 | 81 | $2.9 \pm 0.05$ | 293.00 | 0.150 | 7.5 | (Martin et al., 2016; Jin and Gräter, 2021) |
| CTD2 | 83 | $2.614 \pm 0.05$ | 293.00 | 0.120 | 7.5 | (Jin and Gräter, 2021)(Gibbs et al., 2017)] |
| Sic1 | 92 | $3.0 \pm 0.4$ | 293.00 | 0.200 | 7.5 | (Gomes et al., 2020) |
| SH4UD | 95 | $2.71 \pm 0.04$ | 293.15 | 0.216 | 8.0 | (Shrestha et al., 2019) |
| CoINT | 98 | $2.8 \pm 0.033$ | 277.00 | 0.433 | 7.6 | (Johnson et al., 2017) |
| p15PAF | 111 | $2.81 \pm 0.1$ | 298.00 | 0.150 | 7.0 | (De Biasio et al., 2014) |
| hNL3cyt | 119 | $3.15 \pm 0.2$ | 293.00 | 0.300 | 8.5 | (Paz et al., 2008) |
| RNaseA | 124 | $3.36 \pm 0.1$ | 298.00 | 0.150 | 7.5 | (Riback et al., 2017) |
| +4D | 137 | $2.72 \pm 0.03$ | 298.00 | 0.150 | 7.0 | (Bremer et al., 2022) |
| -3R+3K | 137 | $2.63 \pm 0.02$ | 298.00 | 0.150 | 7.0 | (Bremer et al., 2022) |
| -6R+6K | 137 | $2.79 \pm 0.01$ | 298.00 | 0.150 | 7.0 | (Bremer et al., 2022) |
| -10R+10K | 137 | $2.85 \pm 0.01$ | 298.00 | 0.150 | 7.0 | (Bremer et al., 2022) |
| -4D | 137 | $2.64 \pm 0.01$ | 298.00 | 0.150 | 7.0 | (Bremer et al., 2022) |
| +2R | 137 | $2.62 \pm 0.02$ | 298.00 | 0.150 | 7.0 | (Bremer et al., 2022) |
| +12D | 137 | $2.8 \pm 0.01$ | 298.00 | 0.150 | 7.0 | (Bremer et al., 2022) |
| +12E | 137 | $2.85 \pm 0.01$ | 298.00 | 0.150 | 7.0 | (Bremer et al., 2022) |
| +7K+12D | 137 | $2.92 \pm 0.01$ | 298.00 | 0.150 | 7.0 | (Bremer et al., 2022) |
| +7R | 137 | $2.71 \pm 0.01$ | 298.00 | 0.150 | 7.0 | (Bremer et al., 2022) |
| -12F+12Y-10R | 137 | $2.61 \pm 0.02$ | 298.00 | 0.150 | 7.0 | (Bremer et al., 2022) |
| -10F+7R+12D | 137 | $2.86 \pm 0.01$ | 298.00 | 0.150 | 7.0 | (Bremer et al., 2022) |
| +8D | 137 | $2.69 \pm 0.01$ | 298.00 | 0.150 | 7.0 | (Bremer et al., 2022) |
| +7K+12Db | 137 | $2.56 \pm 0.01$ | 298.00 | 0.150 | 7.0 | (Bremer et al., 2022) |
| -9F+6Y | 137 | $2.66 \pm 0.01$ | 298.00 | 0.150 | 7.0 | (Bremer et al., 2022) |
| -10R | 137 | $2.67 \pm 0.01$ | 298.00 | 0.150 | 7.0 | (Bremer et al., 2022) |
| -9F+3Y | 137 | $2.68 \pm 0.01$ | 298.00 | 0.150 | 7.0 | (Bremer et al., 2022) |
| -8F+4Y | 137 | $2.71 \pm 0.01$ | 298.00 | 0.150 | 7.0 | (Bremer et al., 2022) |
| +7F-7Y | 137 | $2.72 \pm 0.01$ | 298.00 | 0.150 | 7.0 | (Bremer et al., 2022) |
| -12F+12Y | 137 | $2.6 \pm 0.02$ | 298.00 | 0.150 | 7.0 | (Bremer et al., 2022) |
| A1 | 137 | $2.76 \pm 0.02$ | 298.00 | 0.150 | 7.0 | (Bremer et al., 2022) |
| -6R | 137 | $2.57 \pm 0.01$ | 298.00 | 0.150 | 7.0 | (Bremer et al., 2022) |
| aSyn140 | 140 | $3.55 \pm 0.1$ | 293.00 | 0.200 | 7.4 | (Ahmed et al., 2021) |
| FhuA | 144 | $3.34 \pm 0.1$ | 298.00 | 0.150 | 7.5 | (Riback et al., 2017) |
| K27 | 167 | $3.7 \pm 0.2$ | 288.00 | 0.150 | 7.4 | (Mylonas et al., 2008) |
| K10 | 168 | $4.0 \pm 0.1$ | 288.00 | 0.150 | 7.4 | (Mylonas et al., 2008) |
| K25 | 185 | $4.1 \pm 0.2$ | 288.00 | 0.150 | 7.4 | (Mylonas et al., 2008) |
| K32 | 198 | $4.2 \pm 0.3$ | 288.00 | 0.150 | 7.4 | (Mylonas et al., 2008) |
| CAHSD | 227 | $4.84 \pm 0.2$ | 293.00 | 0.070 | 7.0 | (Hesgrove et al., 2021) |
| K23 | 254 | $4.9 \pm 0.2$ | 288.00 | 0.150 | 7.4 | (Mylonas et al., 2008) |
| tau35 | 255 | $4.64 \pm 0.1$ | 293.20 | 0.150 | 7.4 | (Lyu et al., 2021) |
| CoRNID | 271 | $4.7 \pm 0.2$ | 293.15 | 0.192 | 7.5 | (Cordeiro et al., 2019) |
| K44 | 283 | $5.2 \pm 0.2$ | 288.00 | 0.150 | 7.4 | (Mylonas et al., 2008) |
| PNt | 334 | $5.11 \pm 0.1$ | 298.00 | 0.150 | 7.5 | (Riback et al., 2017; Bowman et al., 2020) |
| PNtS1 | 334 | $4.92 \pm 0.1$ | 298.00 | 0.150 | 7.5 | (Bowman et al., 2020) |
| PNtS4 | 334 | $5.34 \pm 0.1$ | 298.00 | 0.150 | 7.5 | (Bowman et al., 2020) |
| PNtS5 | 334 | $4.87 \pm 0.1$ | 298.00 | 0.150 | 7.5 | (Bowman et al., 2020) |
| PNtS6 | 334 | $5.26 \pm 0.1$ | 298.00 | 0.150 | 7.5 | (Bowman et al., 2020) |
| GHRICD | 351 | $6.0 \pm 0.5$ | 298.00 | 0.350 | 7.3 | (Seiffert et al., 2020; Pesce et al., 2023) |

**Table S2.** Experimental solution conditions and PRE data included in the training set for the Bayesian parameter-learning procedure.

| Proteins | N | $N_{\text{labels}}$ | $\omega_I/2\pi$ [MHz] | T [K] | $c_s$ [M] | pH | Ref. |
| --- | --- | --- | --- | --- | --- | --- | --- |
| A2 | 155 | 2 | 850 | 298 | 0.005 | 5.5 | (Ryan et al., 2018) |
| aSyn | 140 | 5 | 700 | 283 | 0.200 | 7.4 | (Dedmon et al., 2005) |
| OPN | 220 | 10 | 800 | 298 | 0.150 | 6.5 | (Kurzbach et al., 2016) |
| FUS | 163 | 3 | 850 | 298 | 0.150 | 5.5 | (Monahan et al., 2017) |
| FUS12E | 164 | 3 | 850 | 298 | 0.150 | 5.5 | (Monahan et al., 2017) |

**Table S3.** Experimental solution conditions and radii of gyration of MDPs included in the training set for the Bayesian parameter-learning procedure.

| Protein | N | $R_g \pm \text{Err}$ [nm] | T [K] | $c_s$ [M] | pH | Ref. |
| --- | --- | --- | --- | --- | --- | --- |
| THB_C2 | 137 | $1.91 \pm 0.076$ | 295.15 | 0.15 | 6.5 | (Michie et al., 2016) |
| Ubq2 | 162 | $2.19 \pm 0.18$ | 293.00 | 0.33 | 8.0 | (Jussupow et al., 2020) |
| Ubq3 | 228 | $2.62 \pm 0.018$ | 293.00 | 0.33 | 8.0 | (Jussupow et al., 2020) |
| Gal3 | 250 | $2.91 \pm 0.06$ | 303.00 | 0.04 | 7.0 | (Lin et al., 2017) |
| TIA1 | 275 | $2.75 \pm 0.05$ | 293.15 | 0.10 | 6.0 | (Sonntag et al., 2017) |
| Ubq4 | 304 | $3.19 \pm 0.092$ | 293.00 | 0.33 | 8.0 | (Jussupow et al., 2020) |
| hnRNPA1* | 314 | $3.12 \pm 0.078$ | 293.15 | 0.15 | 7.5 | (Martin et al., 2021b) |
| hSUMO_hnRNPA1* | 433 | $3.37 \pm 0.13$ | 293.15 | 0.10 | 7.5 | (Martin et al., 2021b) |
| GS0 | 470 | $3.2 \pm 0.044$ | 293.15 | 0.15 | 7.4 | (Moses et al., 2024) |
| GS8 | 486 | $3.37 \pm 0.036$ | 293.15 | 0.15 | 7.4 | (Moses et al., 2024) |
| GS16 | 502 | $3.45 \pm 0.06$ | 293.15 | 0.15 | 7.4 | (Moses et al., 2024) |
| GS24 | 518 | $3.57 \pm 0.075$ | 293.15 | 0.15 | 7.4 | (Moses et al., 2024) |
| GS32 | 534 | $3.75 \pm 0.097$ | 293.15 | 0.15 | 7.4 | (Moses et al., 2024) |
| GS48 | 566 | $4.11 \pm 0.21$ | 293.15 | 0.15 | 7.4 | (Moses et al., 2024) |

**Table S4.**
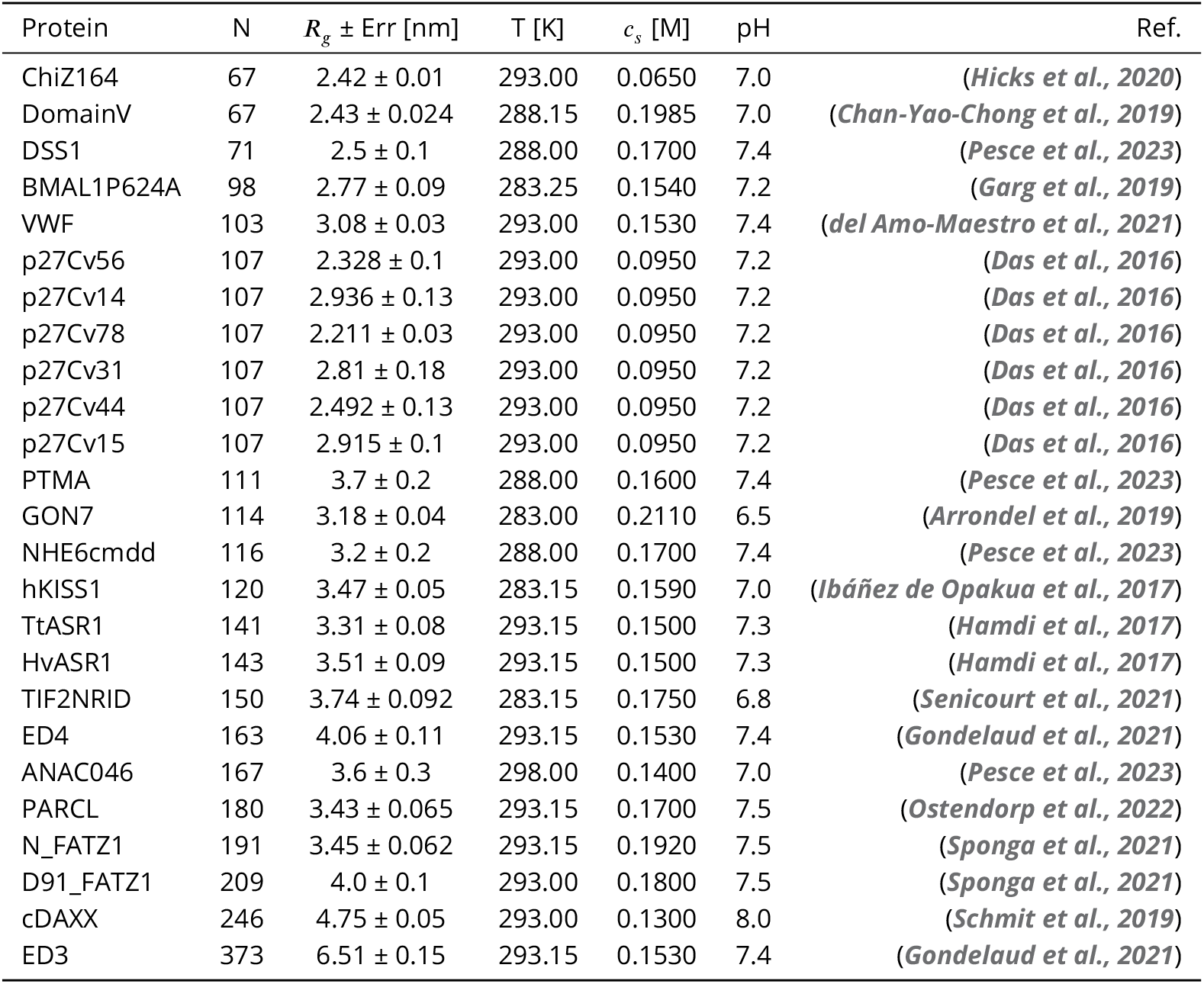
Experimental solution conditions and radii of gyration of IDPs included in the validation set.

**Table S5.** Experimental solution conditions and radii of gyration of MDPs included in the validation set.

| Protein | N | $R_g \pm \text{Err}$ [nm] | T [K] | $c_s$ [M] | pH | Ref. |
| --- | --- | --- | --- | --- | --- | --- |
| SH4UD_SH3_SH2 | 264 | $3.28 \pm 0.06$ | 293.15 | 0.216 | 8.0 | (Gurumoorthy et al., 2023) |
| H46 | 381 | $4.15 \pm 0.05$ | 283.00 | 0.163 | 6.5 | (Elena-Real et al., 2023) |
| TDP43W2A | 415 | $4.11 \pm 0.04$ | 293.15 | 0.312 | 8.0 | (Wright et al., 2020) |
| PCPE | 424 | $4.04 \pm 0.11$ | 293.15 | 0.506 | 7.4 | (Bernocco et al., 2003) |
| NiV_V | 457 | $6.97 \pm 0.02$ | 293.15 | 0.232 | 8.0 | (Salladini et al., 2017) |
| HeV_V | 458 | $6.86 \pm 0.03$ | 293.15 | 0.232 | 8.0 | (Salladini et al., 2017) |
| D14 | 483 | $3.9 \pm 0.17$ | 283.15 | 0.156 | 7.5 | (Hajizadeh et al., 2018) |
| S4FL | 552 | $4.7 \pm 0.1$ | 283.15 | 0.169 | 7.2 | (Gomes et al., 2021) |
| ChiAM | 682 | $4.73 \pm 0.077$ | 293.15 | 0.282 | 8.0 | (Mazurkewich et al., 2020) |

**Table S6.** IDPs and experimental conditions used for slab simulations in this work.

| Proteins | N | $c_s$ [M] | pH | T [K] | $c_{\text{sat,exp}}$ [$\mu\text{M}$] | Ref. |
| --- | --- | --- | --- | --- | --- | --- |
| LAF1 | 176 | 0.15 | 7.5 | 293 | 44.0 | (Schuster et al., 2020) |
| LAF1D2130 | 166 | 0.15 | 7.5 | 293 | 275.0 | (Schuster et al., 2020) |
| LAF1shuf | 176 | 0.15 | 7.5 | 293 | 6.0 | (Schuster et al., 2020) |
| A1S150 | 131 | 0.15 | 7.0 | 293 | 218.1 | (Martin et al., 2021a) |
| A1S200 | 131 | 0.20 | 7.0 | 293 | 159.8 | (Martin et al., 2021a) |
| A1S300 | 131 | 0.30 | 7.0 | 293 | 93.4 | (Martin et al., 2021a) |
| A1S500 | 131 | 0.50 | 7.0 | 293 | 66.5 | (Martin et al., 2021a) |
| -12F+12Y | 137 | 0.15 | 7.0 | 293 | 60.3 | (Bremer et al., 2022) |
| +4D | 137 | 0.15 | 7.0 | 277 | 4.5 | (Bremer et al., 2022) |
| -6R | 137 | 0.15 | 7.0 | 277 | 7.1 | (Bremer et al., 2022) |
| A1 | 137 | 0.15 | 7.0 | 293 | 102.2 | (Bremer et al., 2022) |
| +2R | 137 | 0.15 | 7.0 | 277 | 18.0 | (Bremer et al., 2022) |
| +8D | 137 | 0.15 | 7.0 | 277 | 18.7 | (Bremer et al., 2022) |
| -14N+14Q | 137 | 0.15 | 7.0 | 293 | 171.6 | (Bremer et al., 2022) |
| -10G+10S | 137 | 0.15 | 7.0 | 293 | 268.1 | (Bremer et al., 2022) |
| +7F-7Y | 137 | 0.15 | 7.0 | 293 | 209.0 | (Bremer et al., 2022) |
| -20G+20S | 137 | 0.15 | 7.0 | 293 | 469.4 | (Bremer et al., 2022) |
| -23S+23T | 137 | 0.15 | 7.0 | 293 | 342.2 | (Bremer et al., 2022) |
| -8F+4Y | 137 | 0.15 | 7.0 | 277 | 63.2 | (Bremer et al., 2022) |
| -3R+3K | 137 | 0.15 | 7.0 | 277 | 83.1 | (Bremer et al., 2022) |
| -4D | 137 | 0.15 | 7.0 | 277 | 88.8 | (Bremer et al., 2022) |
| -9F+3Y | 137 | 0.15 | 7.0 | 277 | 115.0 | (Bremer et al., 2022) |
| +23G-23S | 137 | 0.15 | 7.0 | 293 | 46.1 | (Bremer et al., 2022) |
| +23G-23S+7F-7Y | 137 | 0.15 | 7.0 | 293 | 194.0 | (Bremer et al., 2022) |
| +23G-23S-12F+12Y | 137 | 0.15 | 7.0 | 293 | 6.5 | (Bremer et al., 2022) |
| -30G+30S | 137 | 0.15 | 7.0 | 293 | 841.8 | (Bremer et al., 2022) |
| FUS | 163 | 0.15 | 5.5 | 297 | 105.0 | (Murthy et al., 2019) |
| A2 | 155 | 0.01 | 5.5 | 297 | 15.0 | (Ryan et al., 2021) |
| Ddx4WT | 236 | 0.13 | 6.5 | 297 | 230.0 | (Brady et al., 2017) |
| allF | 137 | 0.15 | 7.0 | 293 | 250.0 | (Alshareedah et al., 2023) |
| allY | 137 | 0.15 | 7.0 | 293 | 85.0 | (Alshareedah et al., 2023) |
| allW | 137 | 0.15 | 7.0 | 293 | 1.0 | (Alshareedah et al., 2023) |
| FUS_long | 216 | 0.15 | 7.0 | 285 | 46 | (Farag et al., 2023) |

**Table S7.** Multi-domain proteins and experimental conditions used for slab simulations in this work.

| Proteins | N | $c_s$ [M] | pH | T [K] | $c_{\text{sat,exp}}$ [$\mu\text{M}$] | Box [nm] | Ref. |
| --- | --- | --- | --- | --- | --- | --- | --- |
| hnRNPA1* | 314 | 0.15 | 7.5 | 293 | 173.0 | [20, 20, 270] | (Martin et al., 2021b) |
| hnRNPA3 | 381 | 0.116 | 7.4 | 298 | 6 | [25, 25, 190] | (Kar et al., 2022) |
| hSUMO_hnRNPA1* | 433 | 0.15 | 7.5 | 293 | 136.2 | [25, 25, 190] | (Martin et al., 2021b) |
| FL_FUS | 526 | 0.15 | 7.4 | 293 | 5.5 | [20, 20, 270] | (Wang et al., 2018) |
| GFP_FUS | 764 | 0.15 | 7.4 | 293 | 4.9 | [25, 25, 300] | (Wang et al., 2018) |
| SNAP_FUS | 708 | 0.15 | 7.4 | 293 | 5.9 | [25, 25, 300] | (Wang et al., 2018) |
| SNAP_FUS_PLDY2F_RBDR2K | 710 | 0.15 | 7.4 | 293 | 69.4 | [29, 29, 340] | (Wang et al., 2018) |
| SNAP_FUS_PLDY2F | 710 | 0.15 | 7.4 | 293 | 36.6 | [25, 25, 300] | (Wang et al., 2018) |
| FUS_PLDY2F_RBDR2K | 528 | 0.15 | 7.4 | 293 | 32.0 | [25, 25, 340] | (Wang et al., 2018) |

**Table S8.** Domain boundaries of MDPs used in this study for the Bayesian parameter-learning procedure and validation. Brackets indicate the first and last residue of the domain, respectively. Nested brackets indicate subdomains (restrained) separated by long linkers (unrestrained).

| Protein | N | Domain 1 | Domain 2 | Domain 3 | Domain 4 | Domain 5 | Domain 6 | Domain 7 |
| --- | --- | --- | --- | --- | --- | --- | --- | --- |
| THB_C2 | 137 | [6, 42] | [50, 137] |  |  |  |  |  |
| Ubq2 | 162 | [11, 82] | [87, 158] |  |  |  |  |  |
| Ubq3 | 228 | [1, 72] | [77, 148] | [153, 224] |  |  |  |  |
| Gal3 | 250 | [117, 250] |  |  |  |  |  |  |
| SH4UD_SH3_SH2 | 264 | [94, 150] | [166, 258] |  |  |  |  |  |
| TIA1 | 275 | [6, 82] | [95, 172] | [190, 275] |  |  |  |  |
| Ubq4 | 304 | [1, 72] | [77, 148] | [153, 224] | [229, 300] |  |  |  |
| hnRNPA1* | 314 | [11, 89] | [105, 179] |  |  |  |  |  |
| H46 | 381 | [140, 355] |  |  |  |  |  |  |
| TDP43W2A | 415 | [5, 77] | [107, 177] | [193, 260] | [321, 329] |  |  |  |
| PCPE | 424 | [12, 125] | [134, 249] | [293, 412] |  |  |  |  |
| hSUMO_hnRNPA1* | 433 | [44, 114] | [132, 209] | [224, 298] |  |  |  |  |
| NiV_V | 457 | [406, 457] |  |  |  |  |  |  |
| HeV_V | 458 | [404, 456] |  |  |  |  |  |  |
| GS0 | 470 | [1, 226] | [256, 470] |  |  |  |  |  |
| D14 | 483 | [31, 121] | [157, 246] | [265, 354] | [400, 479] |  |  |  |
| GS8 | 486 | [1, 226] | [272, 486] |  |  |  |  |  |
| hSUMO_TIA1PrLD | 492 | [32, 102] | [114, 186] | [212, 287] | [321, 389] |  |  |  |
| GS16 | 502 | [1, 226] | [288, 502] |  |  |  |  |  |
| GS24 | 518 | [1, 226] | [304, 518] |  |  |  |  |  |
| FL_FUS | 526 | [286, 368] | [423, 451] |  |  |  |  |  |
| FUS_PLDY2F_RBDR2K | 528 | [288, 370] | [425, 453] |  |  |  |  |  |
| GS32 | 534 | [1, 226] | [320, 534] |  |  |  |  |  |
| S4FL | 552 | [15, 138] | [[287, 294],<br>[323, 466],<br>[492, 542]] |  |  |  |  |  |
| GS48 | 566 | [1, 226] | [352, 566] |  |  |  |  |  |
| ChiAM | 682 | [8, 89] | [92, 172] | [178, 257] | [266, 356] | [359, 462] | [471, 567] | [578, 668] |
| SNAP_FUS | 708 | [286, 368] | [423, 451] | [[537, 564], [586, 701]] |  |  |  |  |
| SNAP_FUS_PLDY2F_RBDR2K | 710 | [288, 370] | [425, 453] | [[539, 566], [588, 703]] |  |  |  |  |
| SNAP_FUS_PLDY2F | 710 | [288, 370] | [425, 453] | [[539, 566], [588, 703]] |  |  |  |  |
| GFP_FUS | 764 | [286, 368] | [423, 451] | [529, 755] |  |  |  |  |

**Table S9.** Analysis of the system size effects on slab simulation of hnRNPA1*. The protein concentration is fixed throughout all simulation configurations and is above the experimental saturation concentration. ND: In simulations with 150 chains and box size [11.0, 11.0, 900] nm we did not observe a stable condensed phase.

| | Number of chains & Box size [nm] | Simulation length [μs] | Dilute phase conc. [mM] | Dense phase conc. [mM] |
| --- | --- | --- | --- | --- |
| Varying ($L_x, L_y$) | 45 chains & [11.0, 11.0, 270] | 10 | 0.3±0.1 | 12.0±0.2 |
| | 75 chains & [14.1, 14.1, 270] | 10 | 0.19±0.02 | 12.62±0.02 |
| | 300 chains & [28.3, 28.3, 270] | 10 | 0.18±0.01 | 12.37±0.02 |
| | 450 chains & [34.6, 34.6, 270] | 5 | 0.104±0.008 | 12.32±0.02 |
| Varying $L_z$ | 45 chains & [20.0, 20.0, 81] | 10 | 0.16±0.02 | 11.62±0.05 |
| | 75 chains & [20.0, 20.0, 135] | 10 | 0.15±0.01 | 12.34±0.03 |
| | 300 chains & [20.0, 20.0, 540] | 10 | 0.10±0.01 | 12.51±0.02 |
| | 450 chains & [20.0, 20.0, 810] | 5 | 0.08±0.01 | 12.48±0.03 |
| Varying ($L_x, L_y, L_z$) | 150 chains & [11.0, 11.0, 900] | 10 | ND | ND |
| | 150 chains & [14.1, 14.1, 540] | 10 | 0.17±0.03 | 12.5±0.1 |
| | 150 chains & [28.3, 28.3, 135] | 10 | 0.16±0.01 | 12.11±0.03 |
| | 150 chains & [34.6, 34.6, 90] | 10 | 0.17±0.01 | 10.79±0.06 |

**Table S10.**
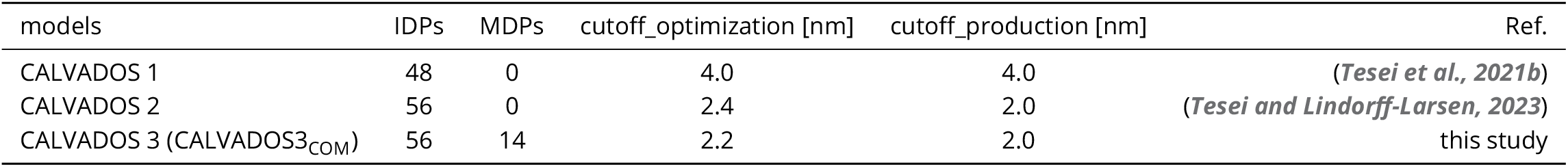
Summary of CALVADOS models. The number of IDPs and MDPs, and cutoff distance for the AH potential used during optimization, cutoff distance of AH potential for validation (production) simulations, and references are shown.

**Figure S1.**
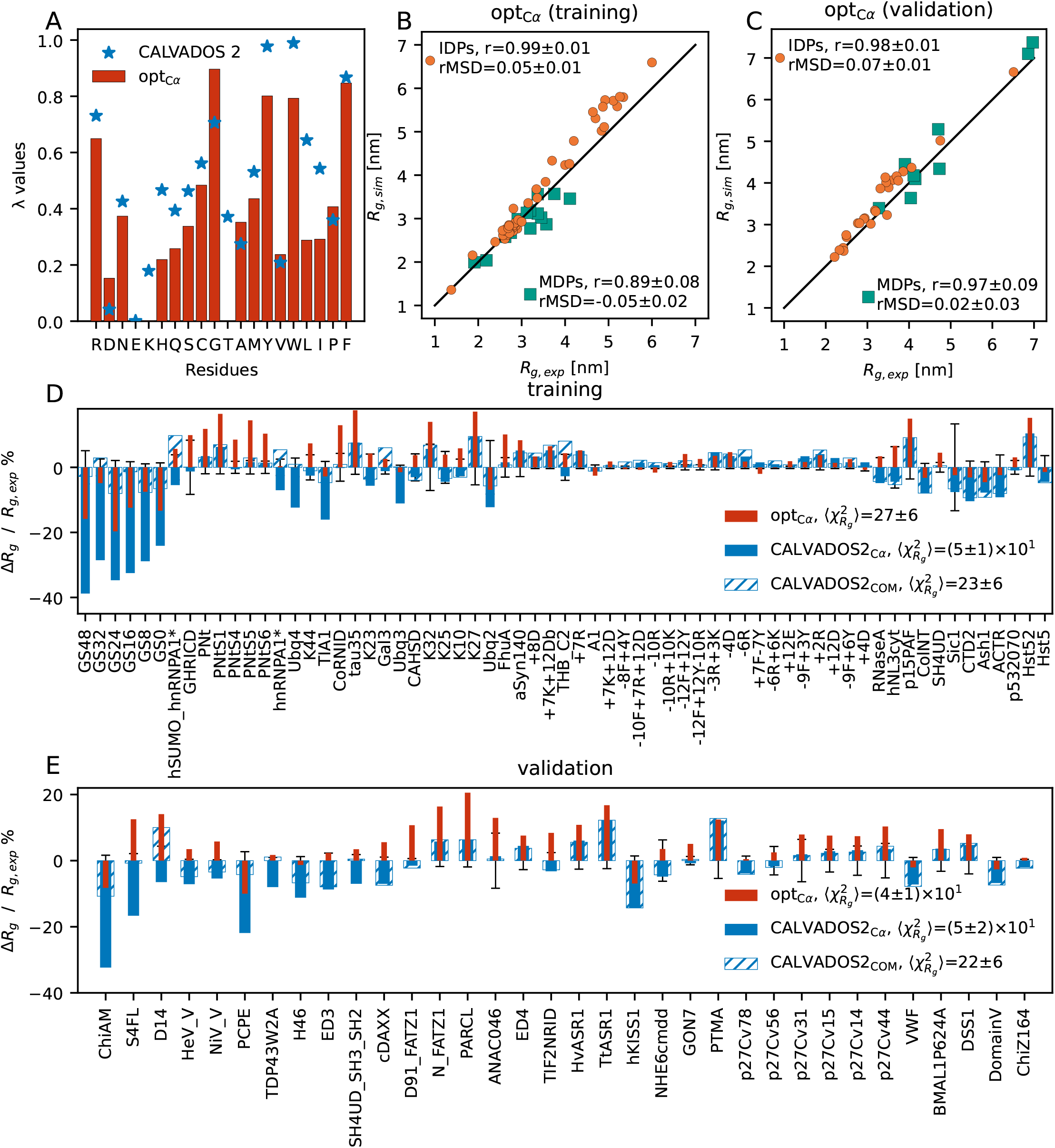
Optimizing the *λ* parameters using a C*α* representation for folded domains. (A) Comparison between *λ* values from CALVADOS 2 (blue) and opt_C*α*_ (red). Comparison between simulated and experimental *R*_*g*_ values for IDPs (orange) and MDPs (green) using opt_C*α*_in (B) the training set and (C) the validation set. Pearson correlation coeicients (*r*) and rMSD are reported in the legend. The black diagonal lines indicate *y* = *x*. Relative difference between experimental and simulated *R*_*g*_ values from opt_C*α*_ (red), CALVADOS2_C*α*_ (blue) and CALVADOS2_COM_ (blue hatched) in (D) the training set and (E) the validation set. 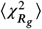 values across IDPs and MDPs are reported in the legend. Error bars show the experimental error divided by *R*_g,exp_. Results from CALVADOS2_C*α*_ and CALVADOS2_COM_ are presented as the same data in ***Figure S4*** and ***Figure 3***.

**Figure S2.**
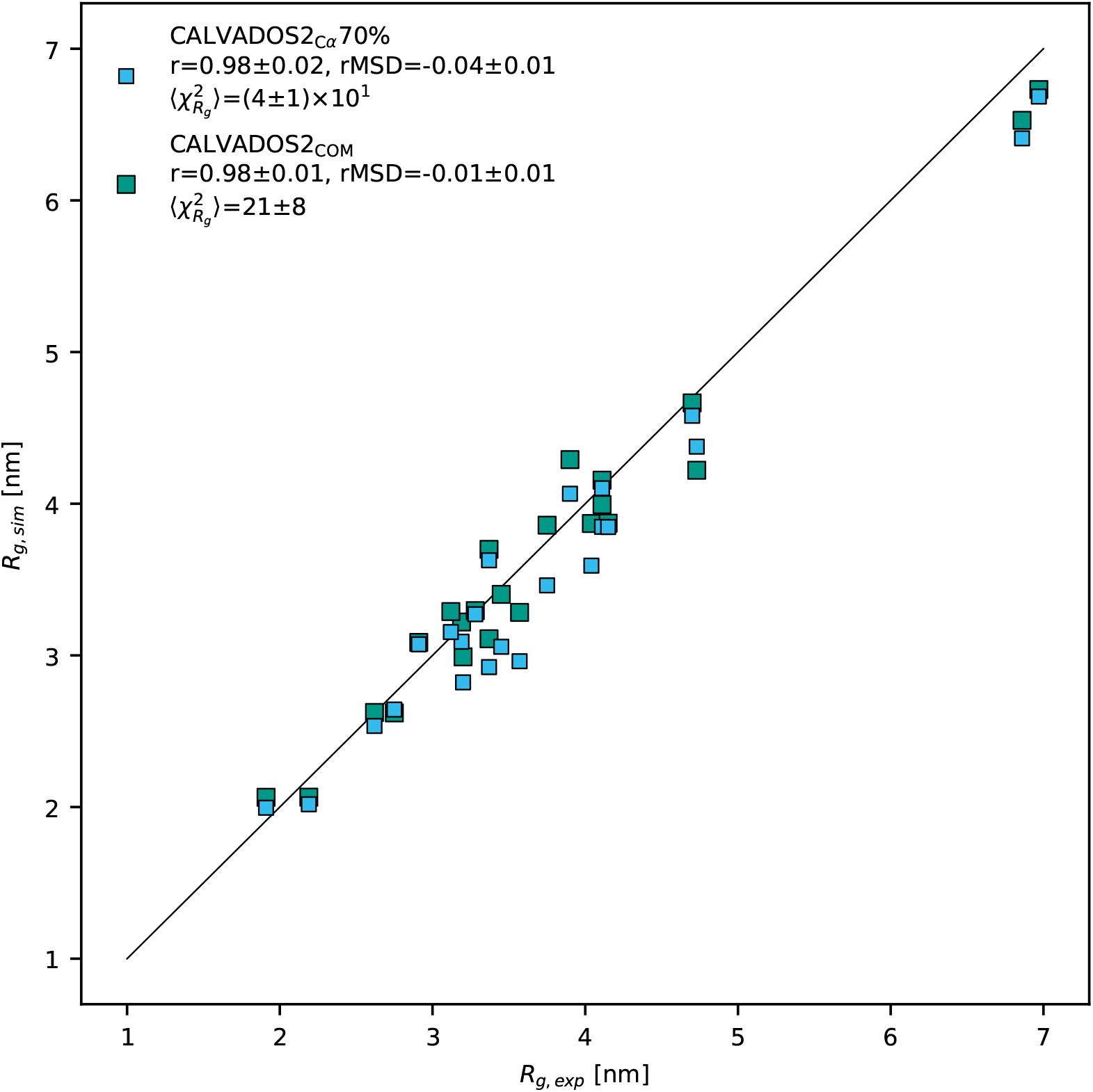
Comparison between simulated and experimental *R*_*g*_ values for all MDPs in the training and validation set using CALVADOS2_C*α*_ 70% (cyan) and CALVADOS2_COM_ (green). Pearson correlation coeicients (*r*), rMSD and 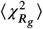 values are reported in the legend. The black diagonal line indicates *y* = *x*.

**Figure S3.**
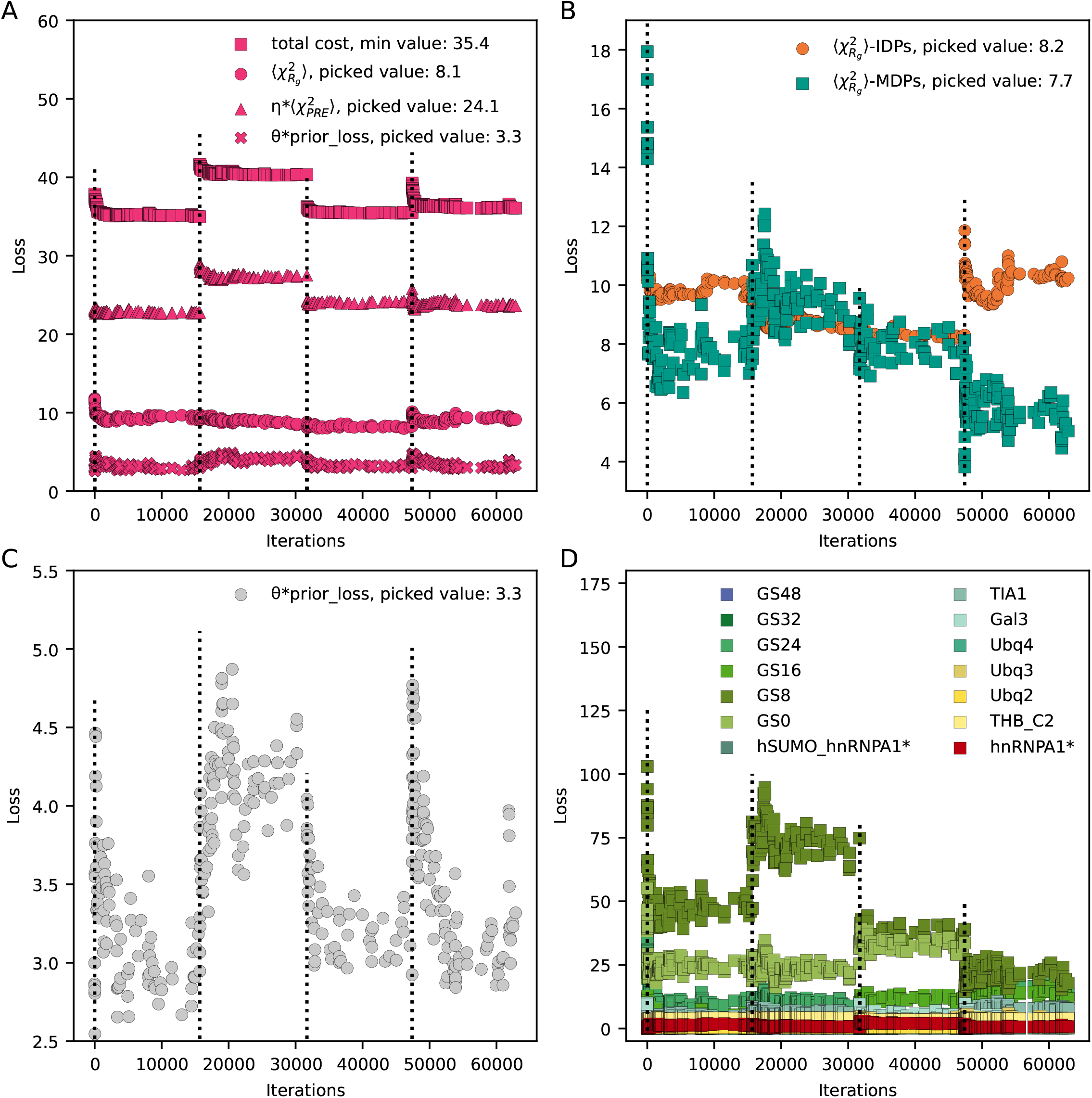
Optimization based on COM representation. The dotted lines indicate the starting points for each cycle. The minimal total cost was used to determine the final *λ* set and the corresponding values of each term are shown in the legends. (A) Evolution of the total cost (square), 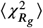 (circle), 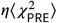 (triangle) and prior loss (cross) in the optimization, with *θ* = 0.08 and *η* = 0.1. (B) Evolution of 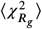 contributed from IDPs (orange) and MDPs (green), respectively. (C) Evolution of prior loss. (D) Evolution of 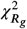 for each MDP during optimization.

**Figure S4.**
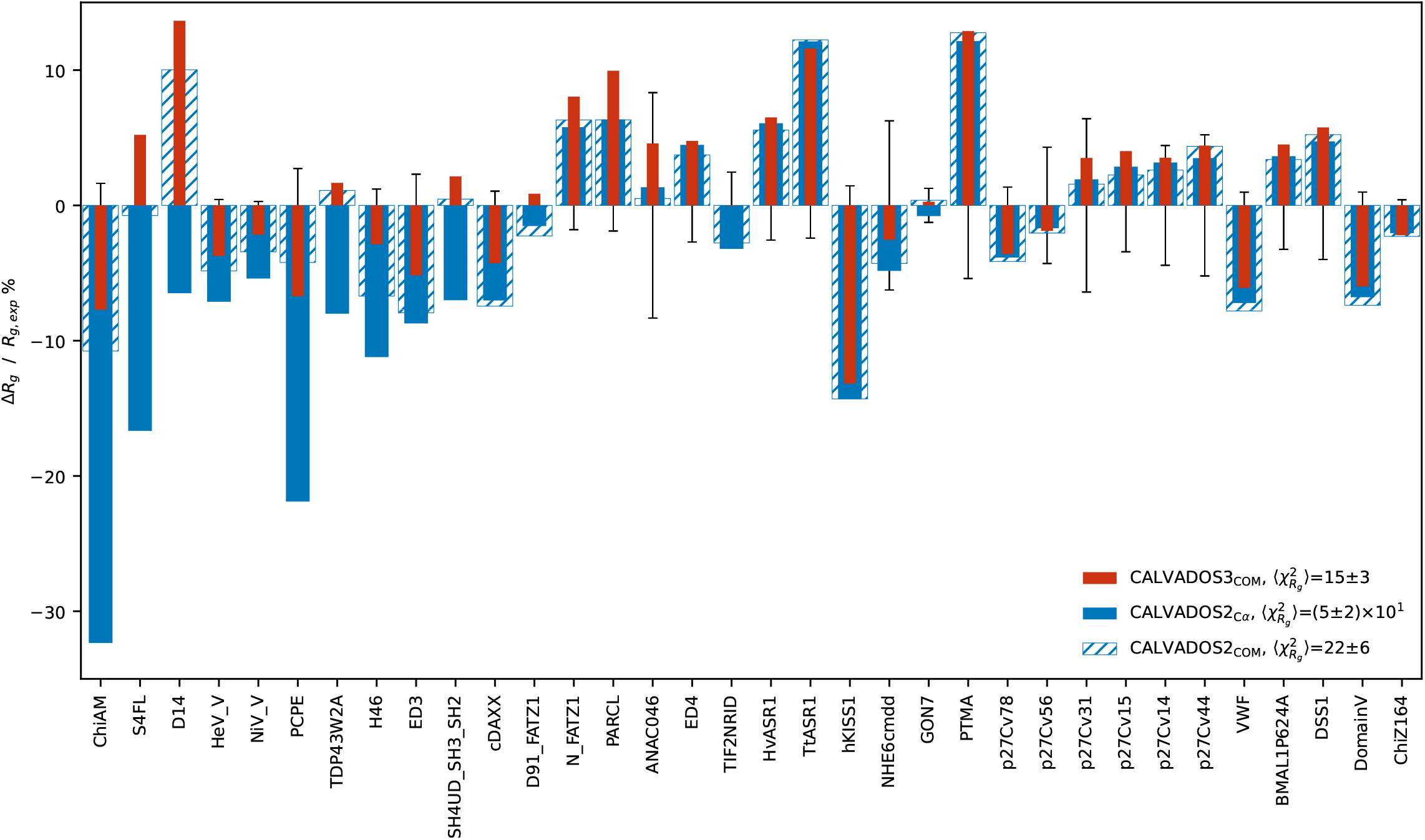
Relative difference between experimental and simulated *R*_*g*_ values from CALVADOS3_COM_ (red), CALVADOS2_C*α*_ (blue) and CALVADOS2_COM_ (blue hatched). 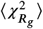 values across IDPs and MDPs in validation set are reported in the legend. Error bars show the experimental error divided by *R* _g,exp_.

**Figure S5.**
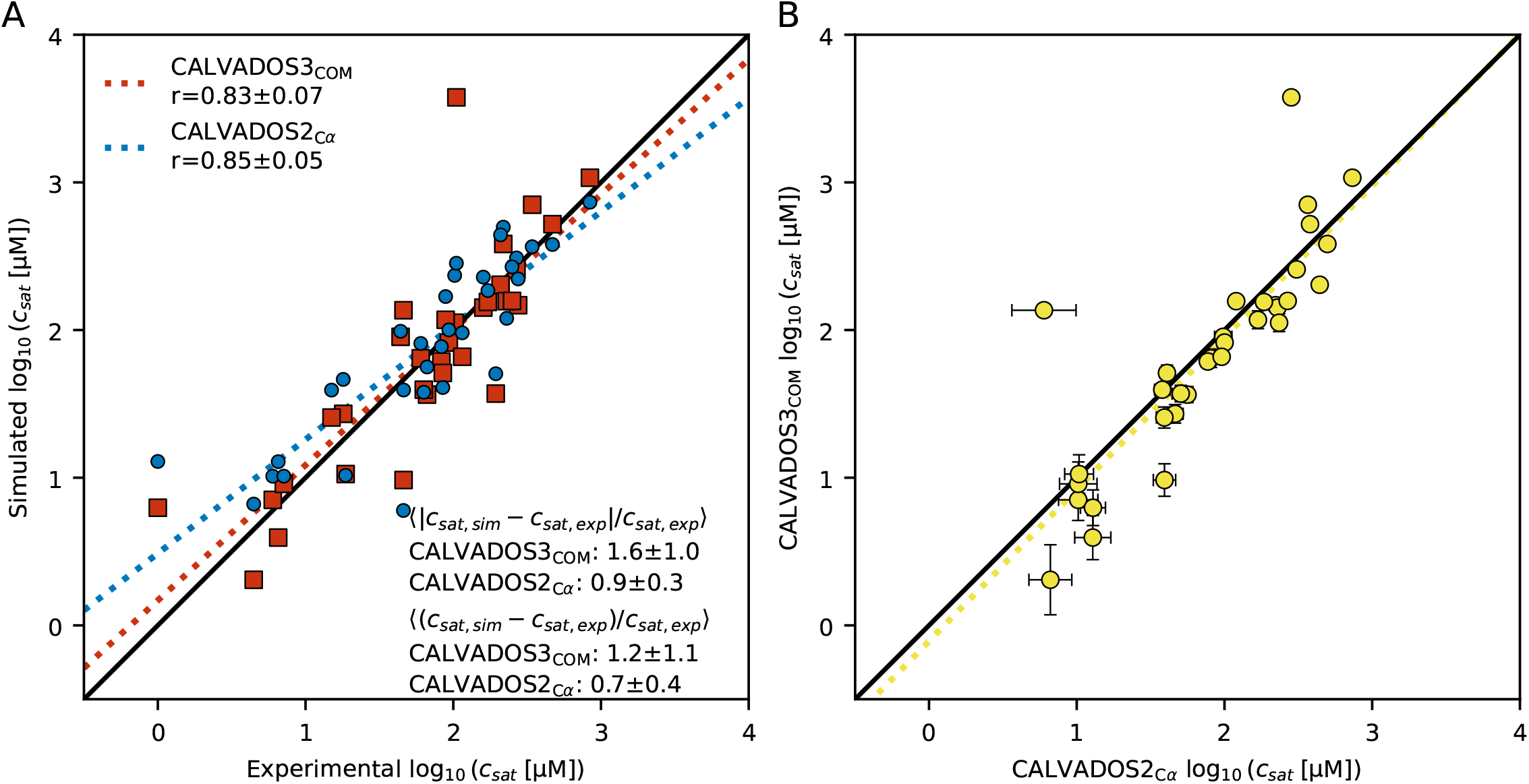
(A) Correlation between *c*_sat_ from simulations yielded by CALVADOS3_COM_ (red) or CALVADOS2_C*α*_ (blue) and experiments for IDPs. (B) Correlation of simulated *c*_sat_ between CALVADOS3_COM_ and CALVADOS2_C*α*_ for the same set of IDPs. Error bars show the error propagation with logarithm from simulated *c*_sat_. The black diagonal lines indicate *y* = *x*. The dotted lines indicate the corresponding linear regression.

**Figure S6.**
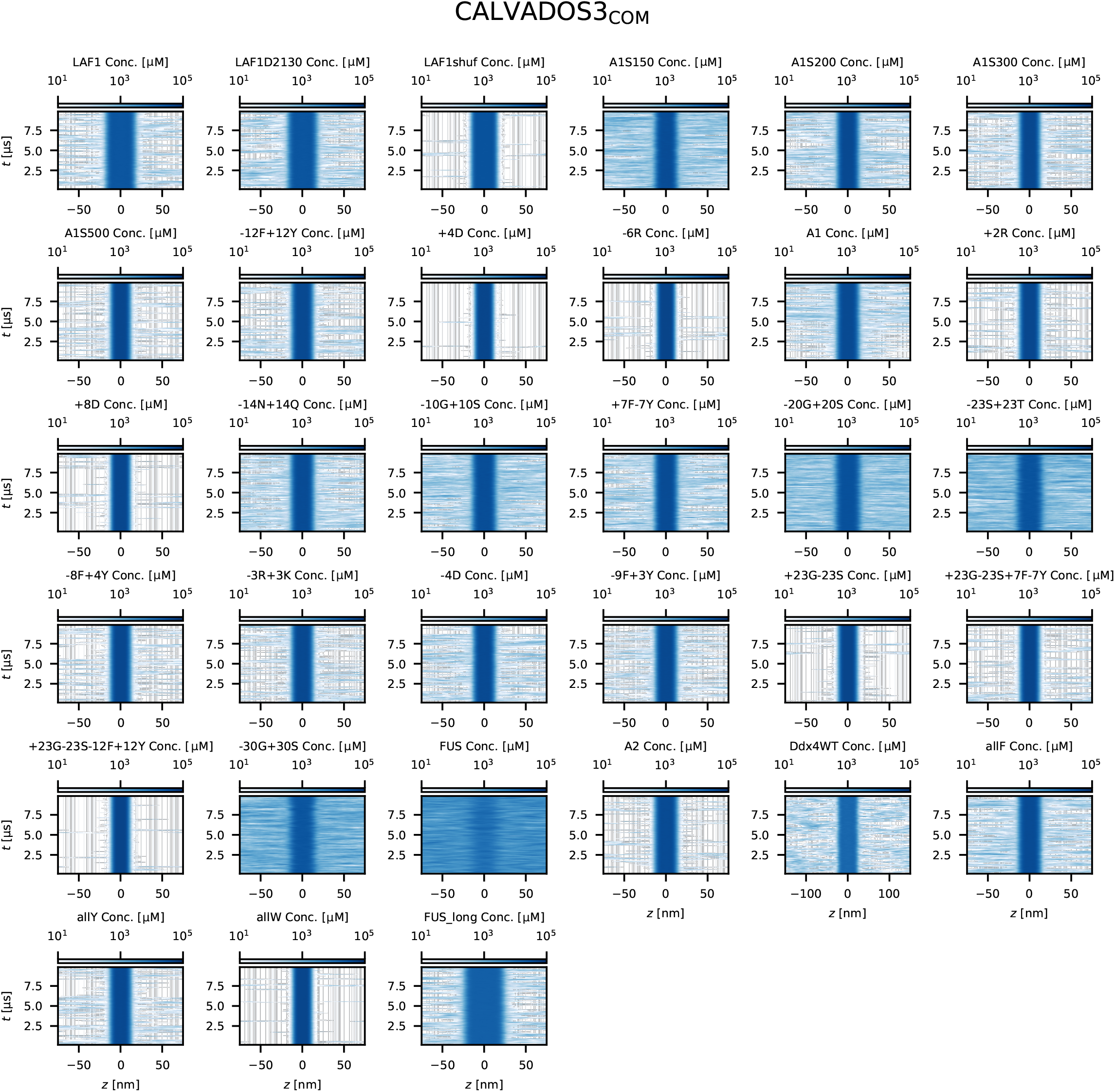
Time evolution of the protein concentration profiles from slab simulations of 33 IDPs using CALVADOS3_COM_ parameters. A more intense colour intensity indicates higher protein concentration.

**Figure S7.**
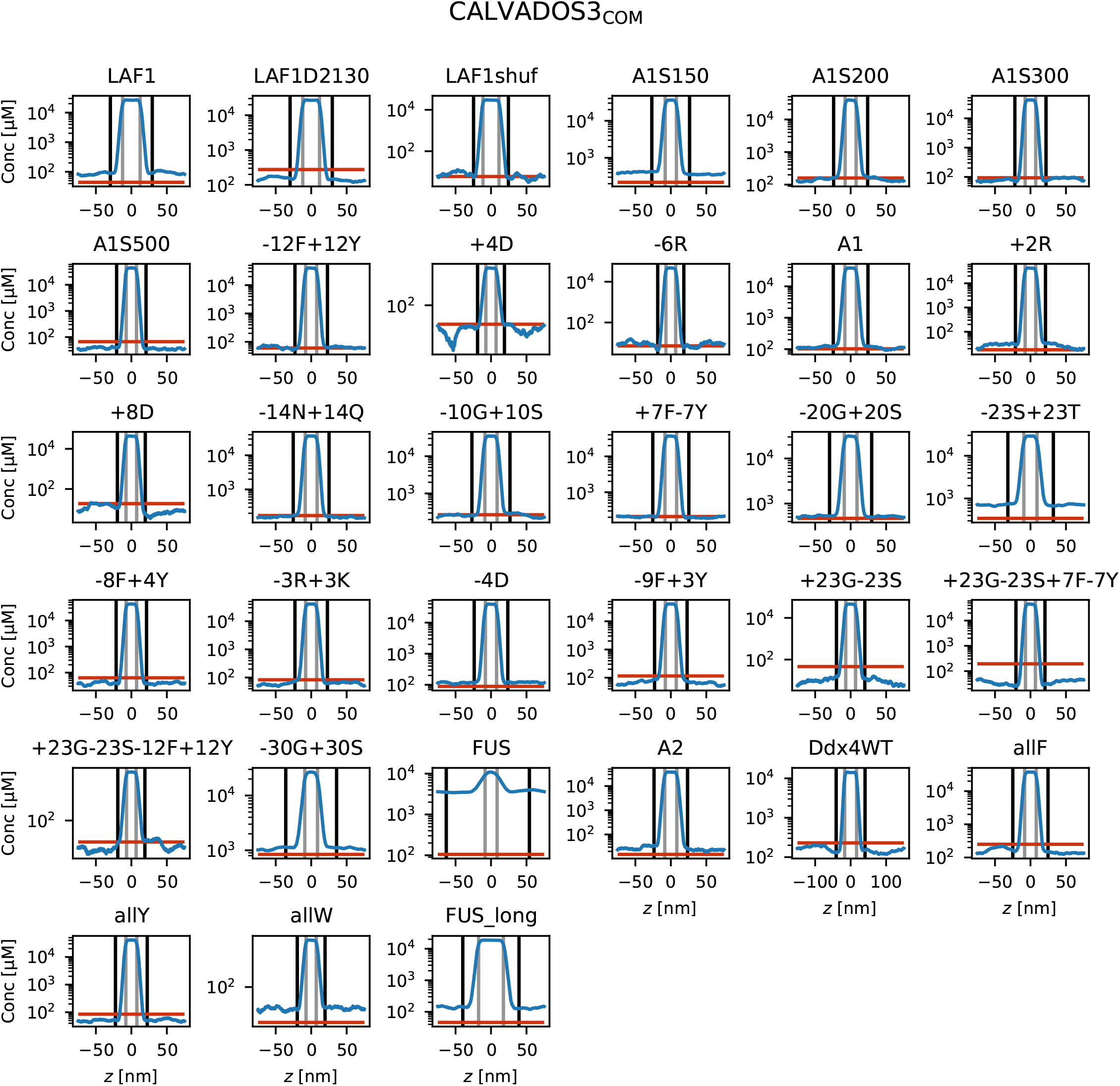
Equilibrium density profiles of slab simulations of 33 IDPs using CALVADOS3_COM_. The red horizontal lines indicate experimental saturation concentrations.

**Figure S8.**
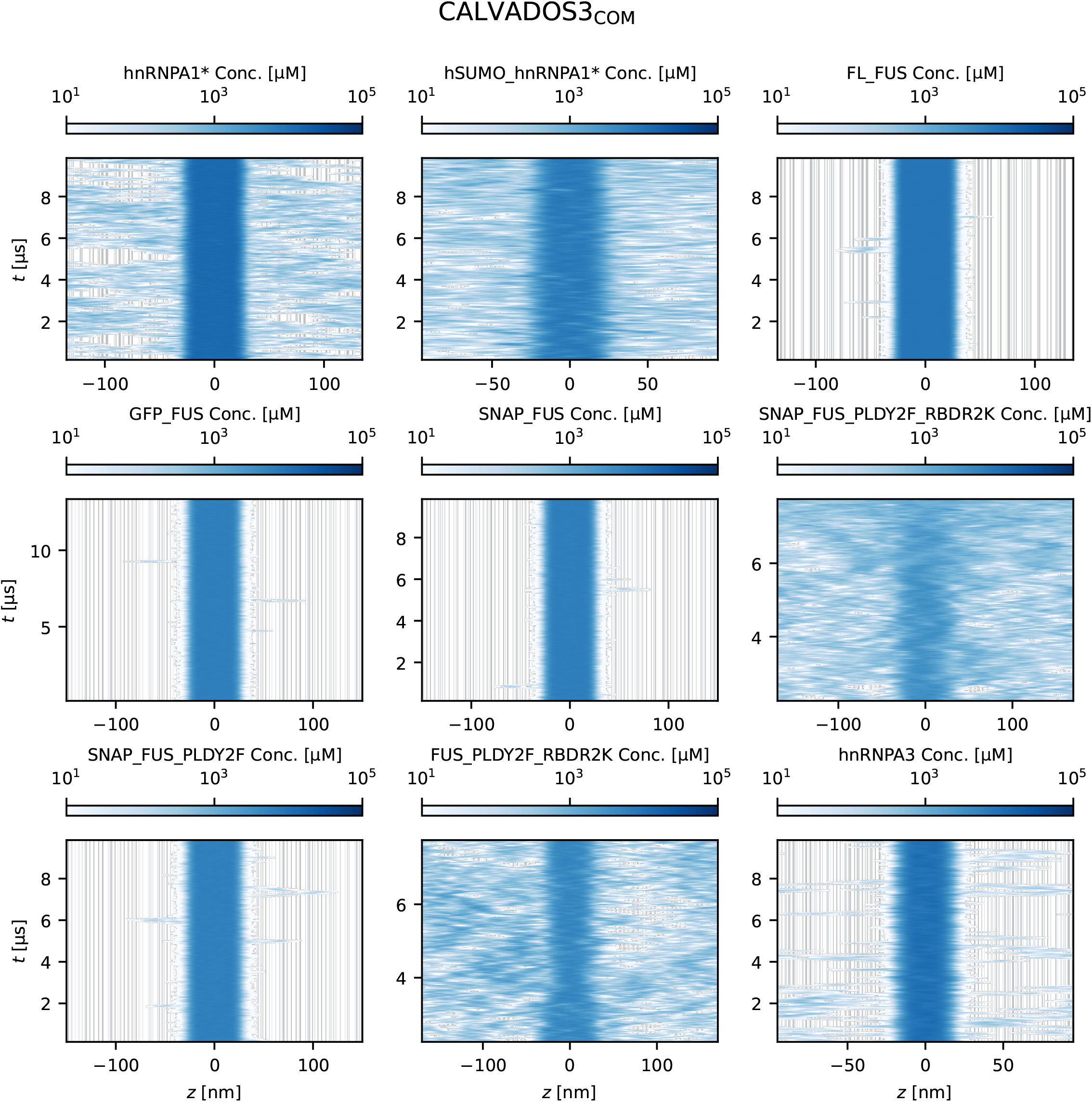
Time evolution of the protein concentration profiles from slab simulations of 9 MDPs using CALVADOS3_COM_ parameters. A more intense colour intensity indicates higher protein concentration.

**Figure S9.**
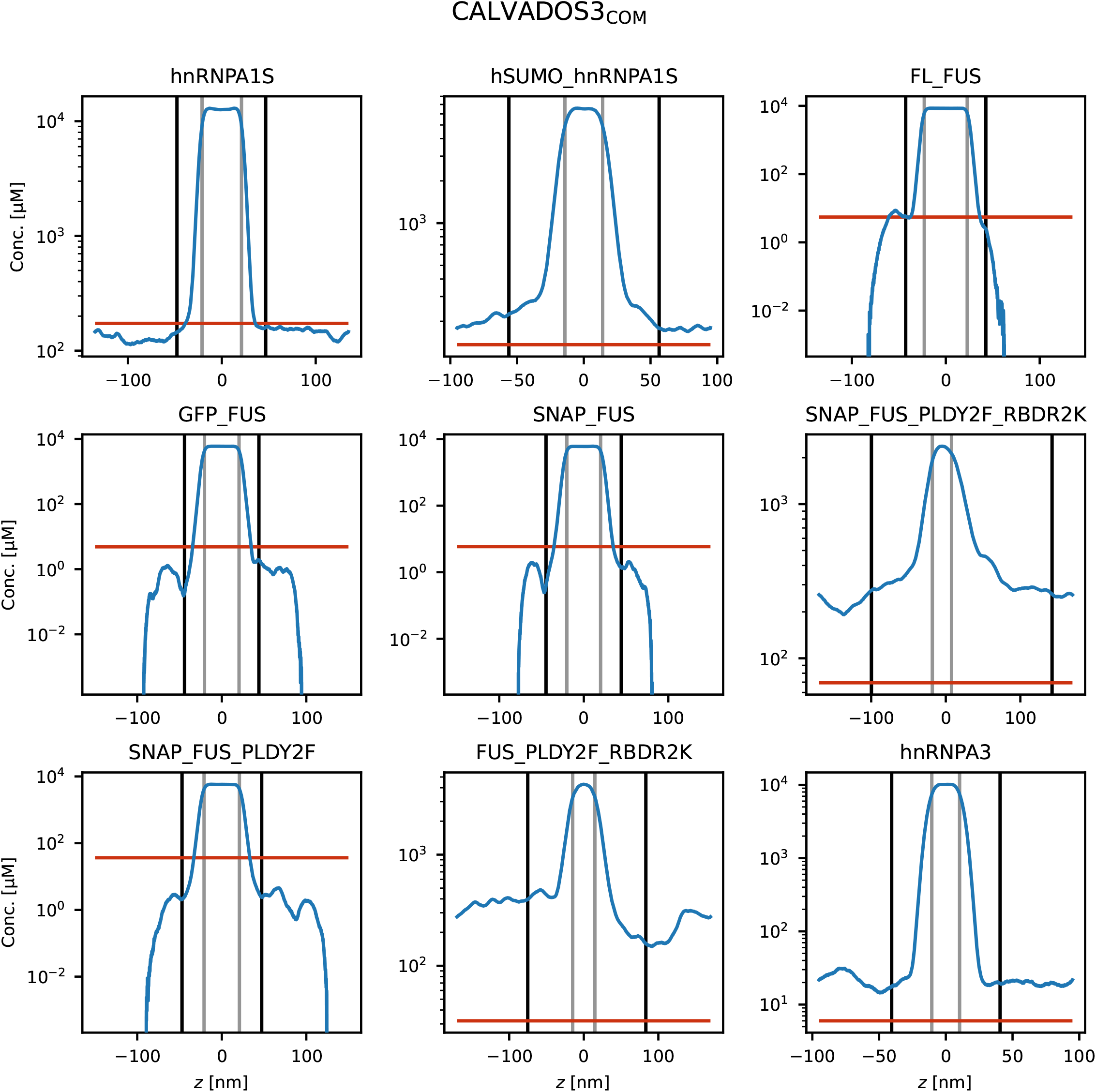
Equilibrium density profiles of slab simulations of nine MDPs using CALVADOS3_COM_. The red horizontal lines indicate experimental saturation concentrations.

**Figure S10.**
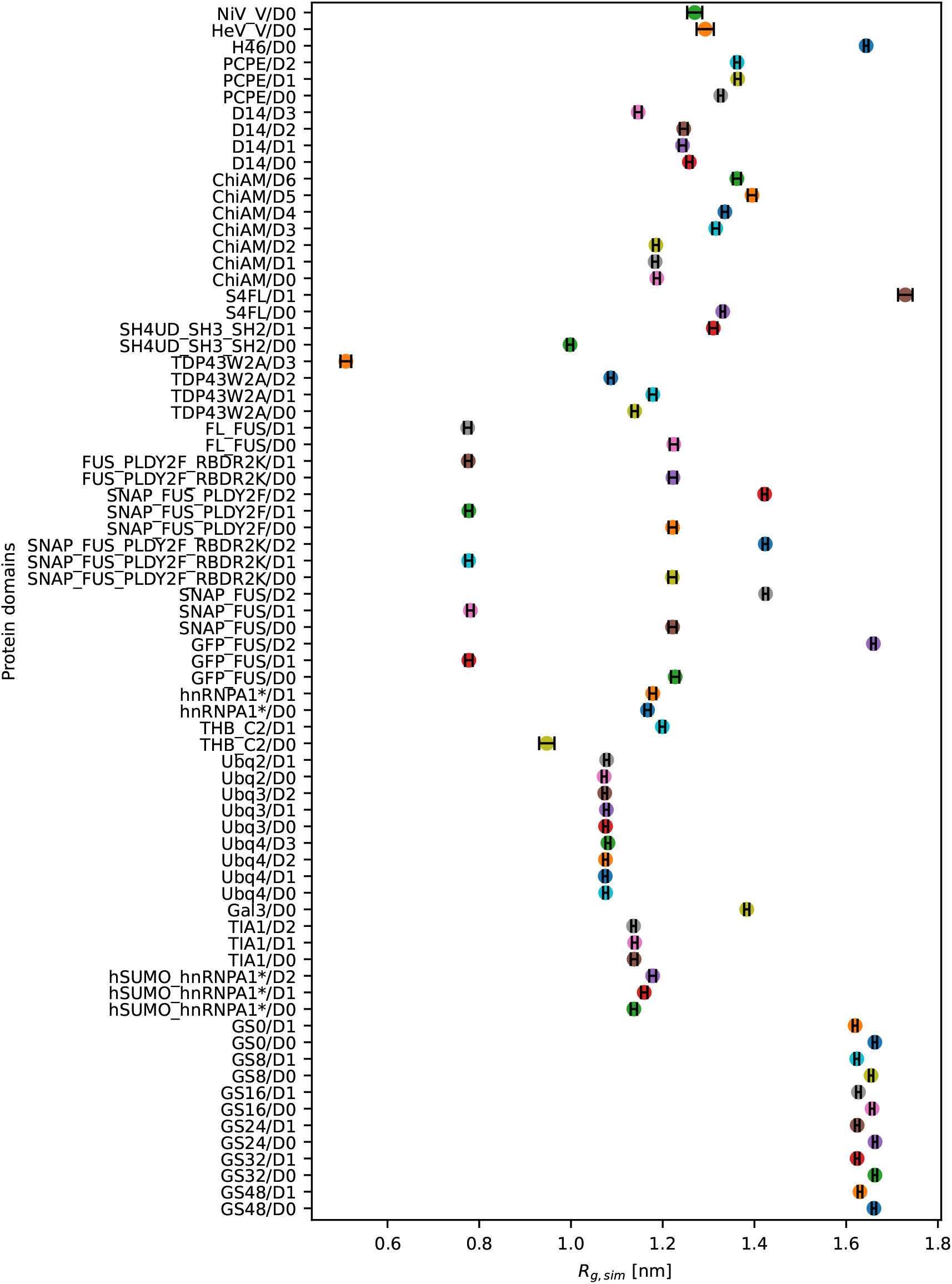
Simulated *R*_*g*_ of domains restrained by elastic network model. Domains in a protein are indicated by D0, D1, D2, etc. Multi-domain proteins in the training set, validation set and slab simulations set are shown.

**Figure S11.**
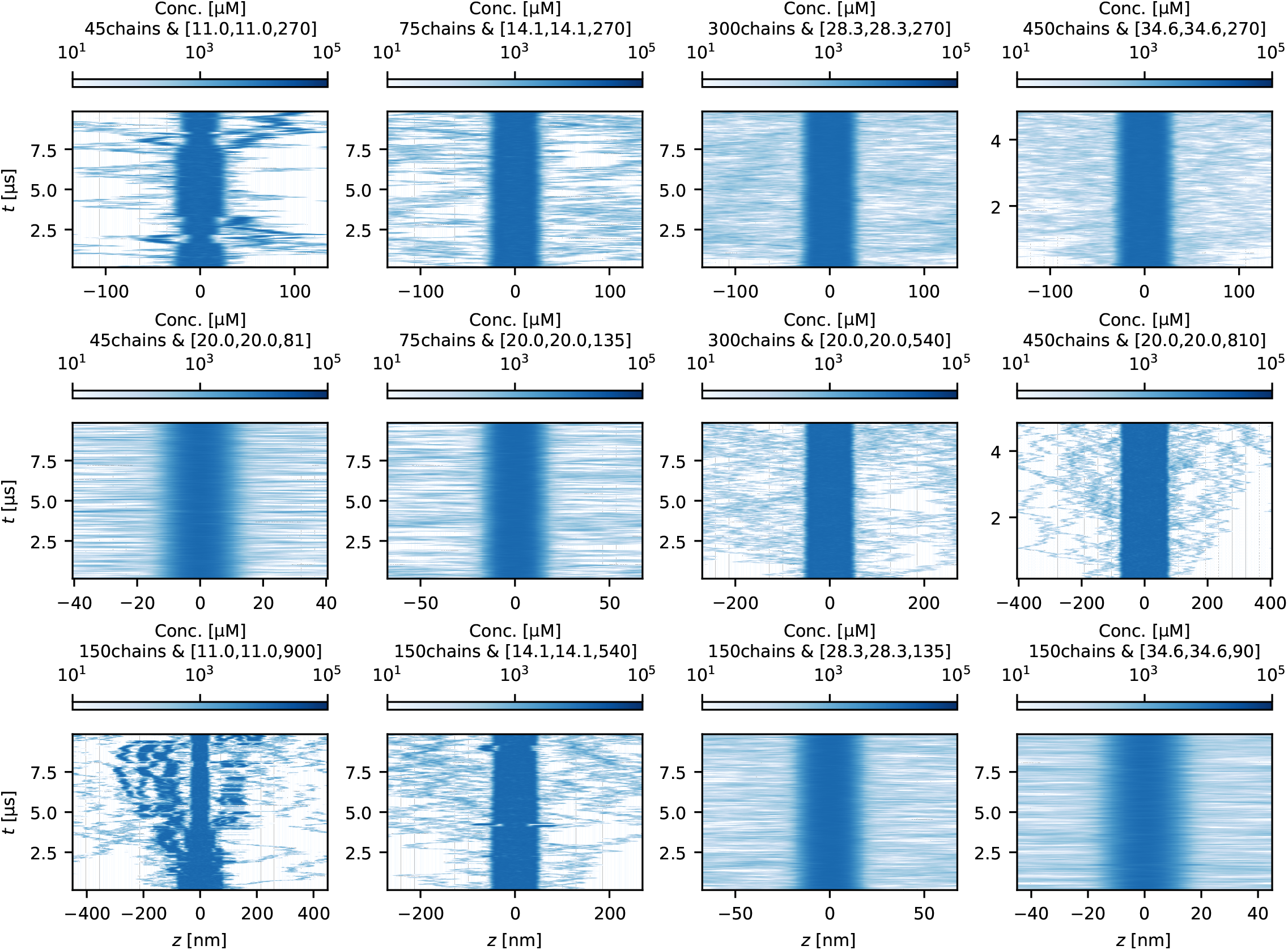
Time evolution of the protein concentration profiles from slab simulations of hnRNPA1* using CALVADOS3_COM_ parameters for analysis of finite-size effects. A more intense colour intensity indicates higher protein concentration. The units of the box sizes are nm.

**Figure S12.**
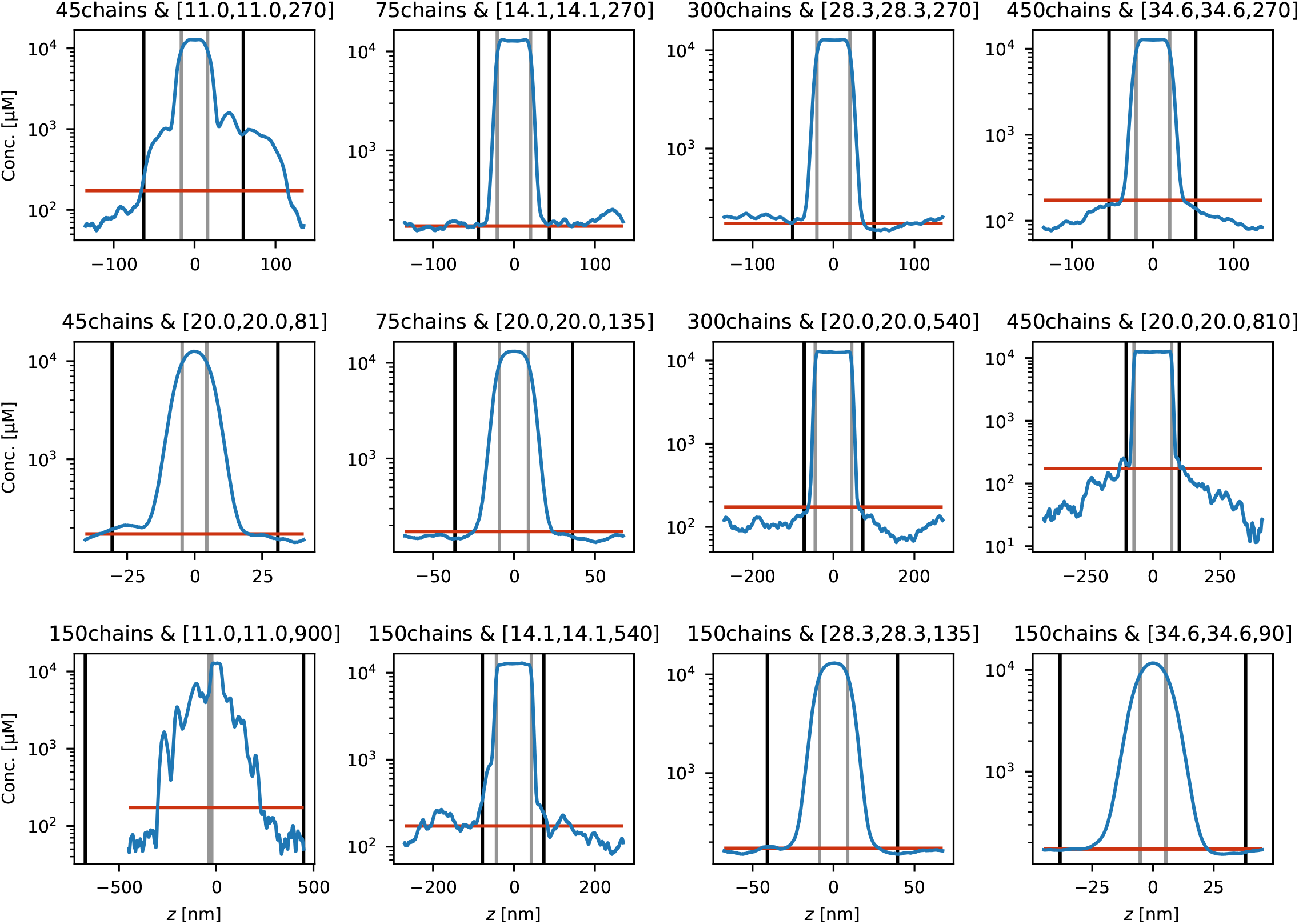
Equilibrium density profiles of slab simulations of hnRNPA1* using CALVADOS3_COM_ for analysis of finite-size effects. The red horizontal lines indicate experimental saturation concentrations. The units of the box sizes are nm.

**Figure S13.**
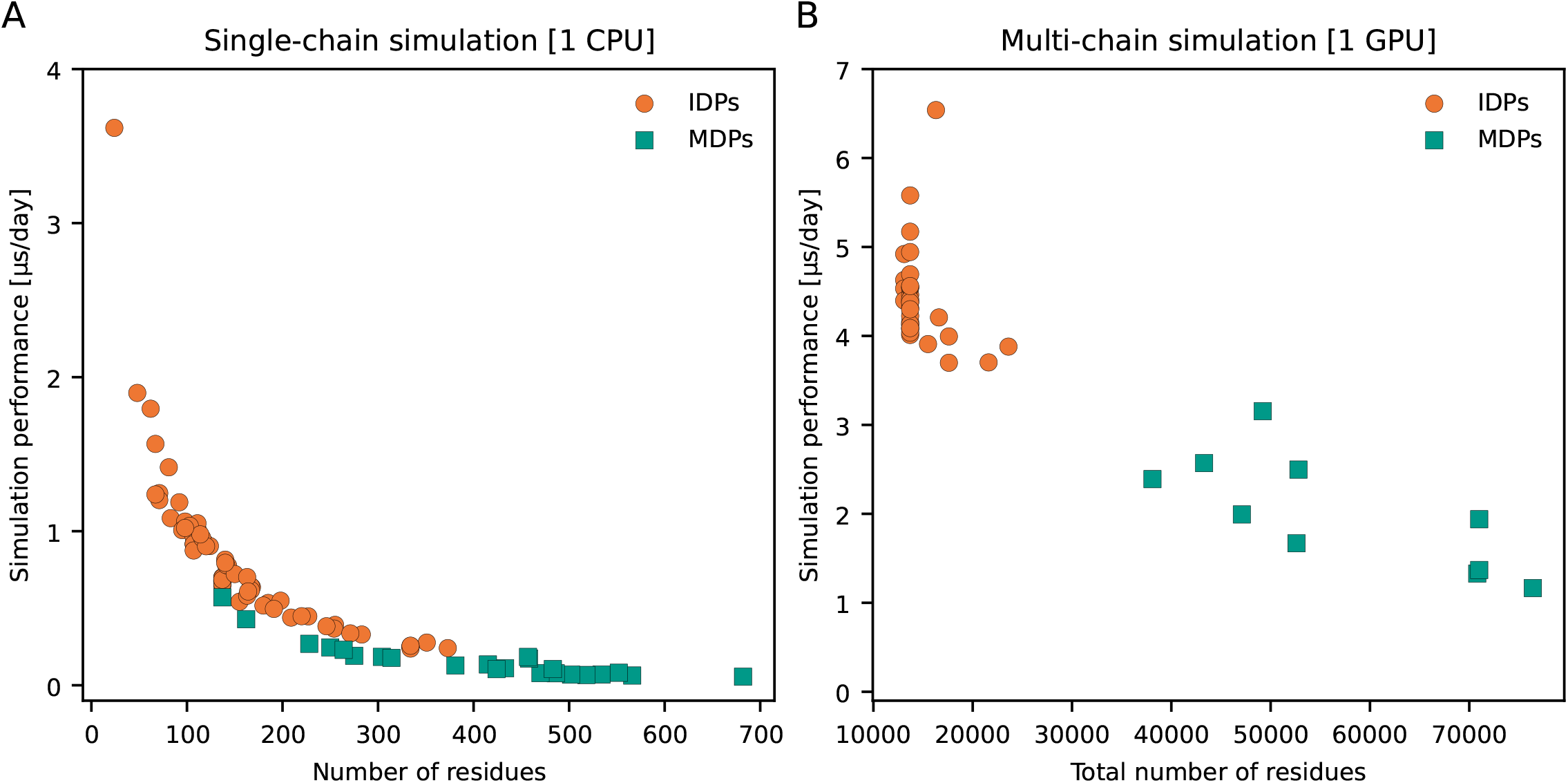
Simulation performance of CALVADOS 3 model on IDPs (orange) and MDPs (green) for (A) single-chain simulations on an Intel Xeon Gold 6130 CPU and (B) multi-chain simulations on an NVIDIA Tesla V100 GPU.

